# Unusual genome expansion and transcription suppression in ectomycorrhizal *Tricholoma matsutake* by repetitive insertions of transposable elements

**DOI:** 10.1101/780601

**Authors:** Byoungnam Min, Hyeokjun Yoon, Julius Park, Youn-Lee Oh, Won-Sik Kong, Jong-Guk Kim, In-Geol Choi

**Author notes:** Corresponding authors (IC); (WK); or (JK). These authors contributed equally to this work (Byoungnam Min, Hyeokjun Yoon).

## Abstract

Genome sequence of *Tricholoma matsutake* was revealed as the one of the large fungal genomes published up to date at 189.0 Mbp with 15,305 predicted genes. The unusual size of this fungal genome contained frequent colonization of transposable elements (TEs) occupying more than half of the entire genome. We identified that 702 genes were surrounded by TEs and 83.2% of those genes were never transcribed at any development stage. This observation corroborated that the insertion of transposable elements alters the transcription of the genes neighboring TEs.

Repeat-induced point mutation such as C to T hypermutation with a bias over ’CpG’ dinucleotides was also recognized in this genome, representing a typical defense mechanism against TEs during evolution. Many transcription factor genes were activated in both primordia and fruiting body, which indicates that many regulatory processes are shared during developmental stages. Small secreted protein genes (<300 aa) were dominantly transcribed in hyphae, where symbiotic interactions occur with hosts. Comparative analysis with 37 Agaricomycetes genomes revealed that IstB-like domain (PF01695) was conserved in taxonomically diverse mycorrhizal genomes, where the *T. matsutake* genome contained four copies of this domain. Three of the IstB-like genes were overexpressed in hyphae. In the CAZyme analysis, reduced CAZyme genes were found as other ectomycorrhizal genomes including a lot of loss of glycoside hydrolase genes. Also, auxiliary activity genes were dominantly transcribed in primordia. The *T. matsutake* genome sequence provides insight into the large genome size and clues to understand unusual fungal genome expansion.

## Introduction

*Tricholoma matsutake* is an ectomycorrhizal (ECM) basidiomycete that establishes a symbiotic relationship with the roots of *Pinus densiflora*, giving it the name “pine mushroom [1].” ECM fungi build an aggregated hyphal sheath that encases the whole root tip of its symbiotic partner and mediates the root’s external interactions with the soil [2]. This encasing or root colonization is formed through a hyphal network called the “Hartig net” that is located inside the root cells, an anatomical pattern shared by a majority of ECM fungi [3, 4]. The fruiting body of *T. matsutake* is a highly valued edible mushroom in many countries [1, 5]. Unfortunately, attempts to cultivate the fruiting body have been unsuccessful and the mechanism for mushroom development not yet fully understood.

Mushroom formation is proceeded by distinct developmental stages that include vegetative hyphae, dikaryotic primordia, and mature fruiting body [6]. Various genes including transcriptional factors [7], hydrophobins [8], and light receptors [9] have been suggested as critical genetic factors for the fruiting body formation in basidiomycetes. Systematic transcriptomic surveys on fruiting body formation have been carried out for various basidiomycetes [10].

Transposable elements (TE) play an important role in genome evolution by causing chromosomal rearrangements or reshaping regulatory networks [11, 12]. Many ECM genomes show high TE content leading to comparably larger genome sizes [13] and contain more TEs than asymbiotic relatives [14]. The effect of TEs in mushrooms is transcriptional repression, particularly when genes are enclosed by TEs [15]. In a recent comparative genomic study of two mushroom strains, *Pleurotus ostreatus* PC15 and PC9, the genes surrounded by transposons in one strain showed strong transcriptional repression, whereas their orthologs in another strain were normally expressed [15]. Despite the higher TE content in ECM genomes, the transcription tendency of the ECM genes affected by TEs has not been thoroughly examined.

Here we report the genome sequence of *T. matsutake* and the transcriptional dynamics over three distinct developmental stages. The most distinct features of *T. matsutake* genome were genome expansion by the many TEs and prevailing transcriptional suppression in all developmental stages. In addition, we performed comparative analyses on the *T. matsutake* and 37 Agaricomycetes genomes to identify potential gene clusters involved in symbiosis.

## Results and Discussion

### Genomic summary of *T. matsutake*

Sequencing of the dikaryotic genomic DNA of *T. matsutake* generated a total length of 189.0 Mbp within 5,255 scaffolds with 111.8× sequencing coverage. We predicted 15,305 gene models using FunGAP pipeline [16]. The predicted genes were examined for their reliability by RNA-seq, functional domains, and orthologs, thereby the 14,528 (94.9%) genes were supported by at least one evidence (S1 Fig). Genome completeness test using BUSCO v3.0.2 [17] showed >99% coverage of single-copy orthologs in Basidiomycota (1,323 of 1,335 entries), validating the complete genome assembly and annotation. Because the genome was dikaryotic, we investigated how many genes are allelic by comparing two-member gene families with relative genomes. As a result, we identified that allele genes were not frequent in the assembled genome because of its lack of two-member gene family expansion (S1 Fig). We also confirmed that there was no contaminated sequence in the final assembly (S2 Fig). K-mer frequency of genomic DNA reads showed a bimodality indicating the diploidy (S3 Fig).

As of Sep 2019, genome sizes of sequenced fungi deposited in the NCBI ranged from 2 Mbp to 2.1 Gbp with an average of 31.0 Mbp (40.7 Mbp for basidiomycetes), and *T. matsutake* genome had relatively large size (Fig 1). Contrary to the size of the genome, the gene-to-genome ratio was comparably low (81 genes per Mbp). This indicates the presence of many noncoding DNA regions (e.g., repetitive elements). Data concerning genome assembly and predicted genes are summarized in Table 1. The species tree of *T. matsutake* with 37 Agaricomycetes is shown in Fig 1.

**Fig 1.**
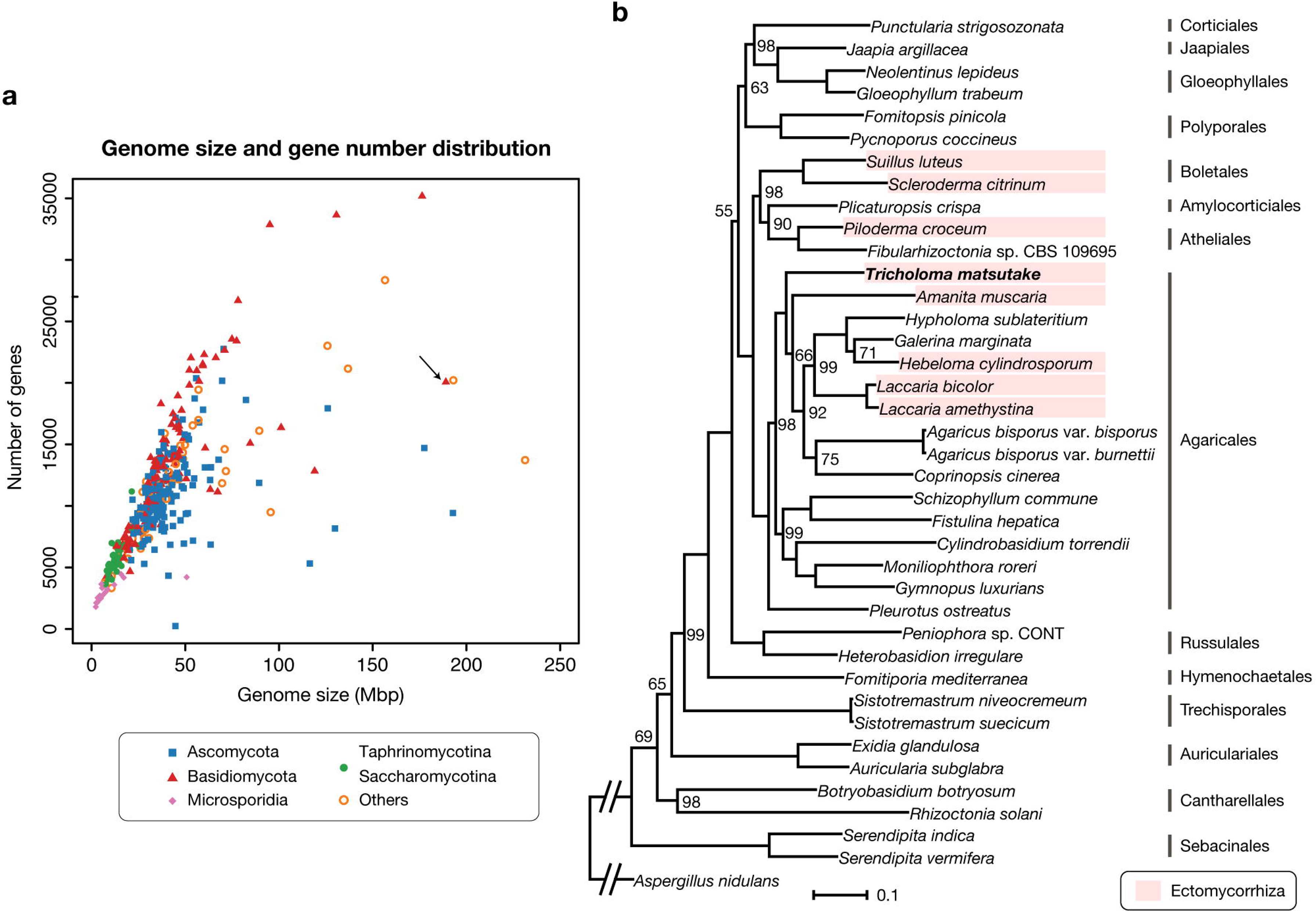
The genome sequencing of *Tricholoma matsutake*. a. Genome size vs. gene number of all available fungal genomes in the NCBI. As of Sep 2019, 5,415 fungal genome assemblies have been deposited and 1,618 had gene predictions. We used one genome per genus to draw the plot. *Tricholoma matsutake* is indicated by the arrow. **b.** Species tree of *Tricholoma matsutake* and 37 Agaricomycetes genomes. Only bootstrap values of less than 100 are marked. The scale bar that represents the mean number of amino acid substitutions per site is shown. *Aspergillus nidulans* genome (GenBank: GCF_000149205.2) was used as an outgroup. The branch to the outgroup was shortened for visualization purpose.

**Table 1.** Genomic features of Tricholoma matsutake

| **Assembly statistics** |  |
| --- | --- |
| Total contig length | 144.2 Mbp |
| Total scaffold length | 189.0 Mbp |
| Average base coverage | 109.8× |
| Number of contigs | 29,547 |
| Number of scaffolds | 5,255 |
| N50 contig length | 10.2 kbp |
| N50 scaffold length | 93.4 kbp |
| G+C content (overall) | 45.11% |
| G+C content (coding region) | 49.57% |
| G+C content (non-coding region) | 44.51% |
| Repeat elements | 92.4 Mbp |
| **Predicted protein-coding genes** |  |
| Predicted genes | 15,305 |
| Percent coding | 8.94% |
| Average coding sequence size | 1,104.04 nt |
| Gene density | 80.97 genes/Mbp |
| Total exons | 81,887 |
| Total introns | 66,582 |
| Number of introns per gene (median) | 4 |
| Number of exons per gene (median) | 4 |
| Average exon length | 206.35 nt |
| Average intron length | 77.14 nt |

### Repeat elements in the *T. matsutake* genome

The *T. matsutake* genome had a high content of repeat elements, which has been suggested as a concurrent feature of ectomycorrhizal genomes [2,13,14]. The repeat elements were estimated to have a total size of 92.4 Mbp, representing 48.9% of the entire 189.0 Mbp genome (Fig 2). The major classes of repeat elements were LTR/Gypsy and LTR/Copia transposable elements, corresponding to 41 Mbp (21.9%) and 7 Mbp (3.9%) of the genome, respectively. These two frequent elements have also been observed in other basidiomycete genomes [14, 15]. Within the repeat regions, there were 15,014 complete coding sequences (containing start codon, stop codon, and no internal stop codon), which was equivalent to a total size of 10.9 Mbp. They were not included in the final predicted genes or further functional annotation. These sequences were annotated as reverse transcriptase (PF00078) (12%), integrase (PF00665) (6%), and chromodomain (PF00385) (5%). A majority of these sequences (14,857 of 15,014, 99.0%) were transcriptionally repressed (zero Fragments Per Kilobase of transcript per Million mapped reads (FPKMs) at all developmental stages). We presumed that meiotic silencing by unpaired DNA and the quelling process are the potential repression mechanisms [18], because the genome included the genes responsible for those processes (*sad-1*, *sms-3*, and *sms-2* for meiotic silencing by unpaired DNA, and *qde-1*, *dcl2*, and *qde2* for the quelling process; Fig 3). Another presumed mechanism is the repeat-induced point mutation (RIP). Despite a previous report that RIP does not exist in Agaricomycotina genomes [19], we identified the pattern of CpG hypermutations in the genome though further studies remain to reveal whether actual RIP process made this pattern (S4 Fig). Although the genome lacked *rid/dim2* responsible for RIP process [20] (Fig 3), its homolog, *masc2*, existed with two copies. This pattern is also frequent in other basidiomycete genomes [21]. Experimental validation remains to be done to reveal the exact function of Masc2 in the control of the RIP process. Intergenic region length distribution indicated that many genes were located in gene-sparse regions mainly due to enriched transposable elements (S5 Fig).

**Fig 2.**
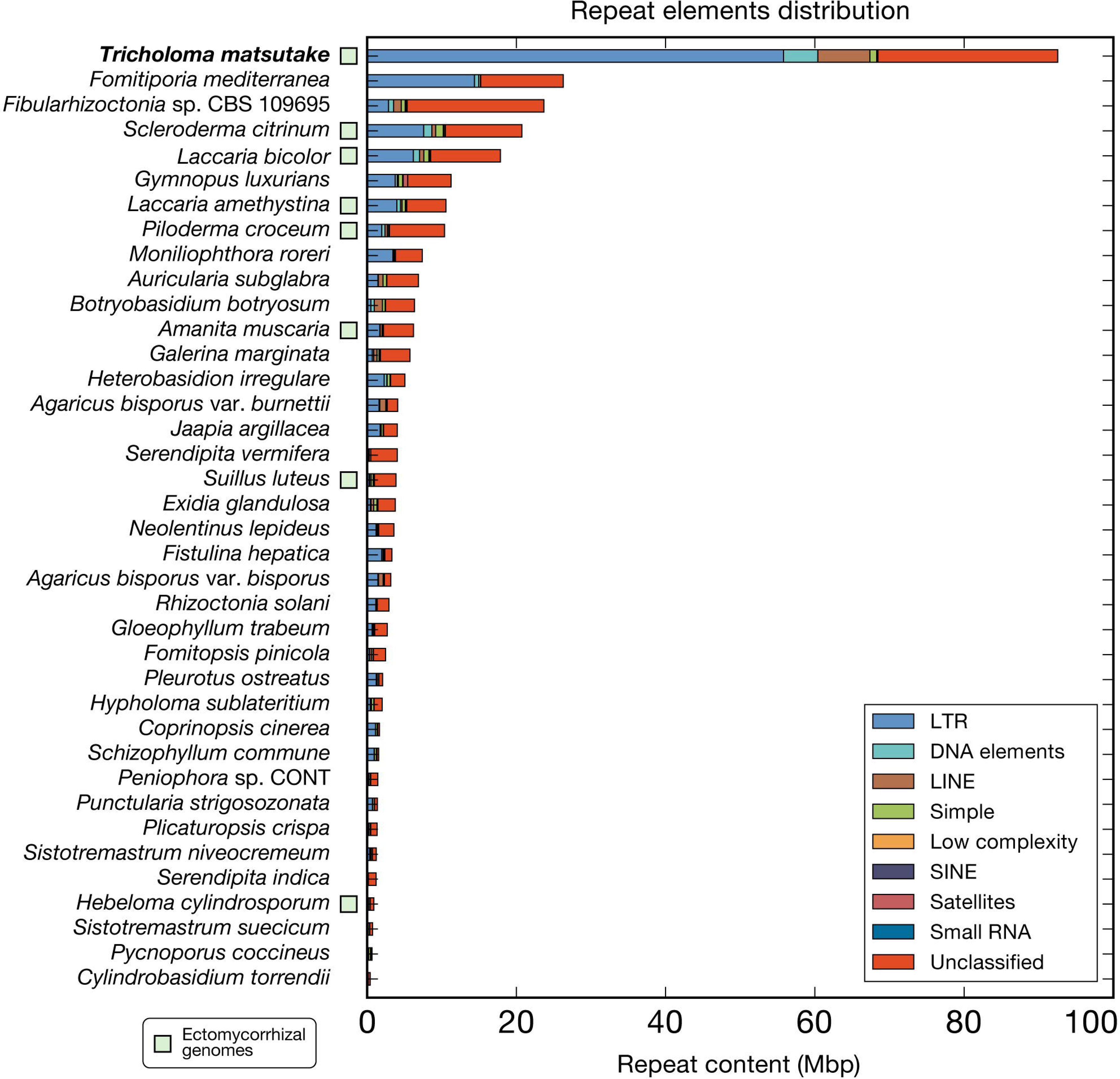
Repeat content in the *Tricholoma matsutake* and 37 Agaricomycetes genomes. RepeatModeler and RepeatMasker (http://www.repeatmasker.org) were sequentially used to predict repeat elements in the genomes.

**Fig 3.**
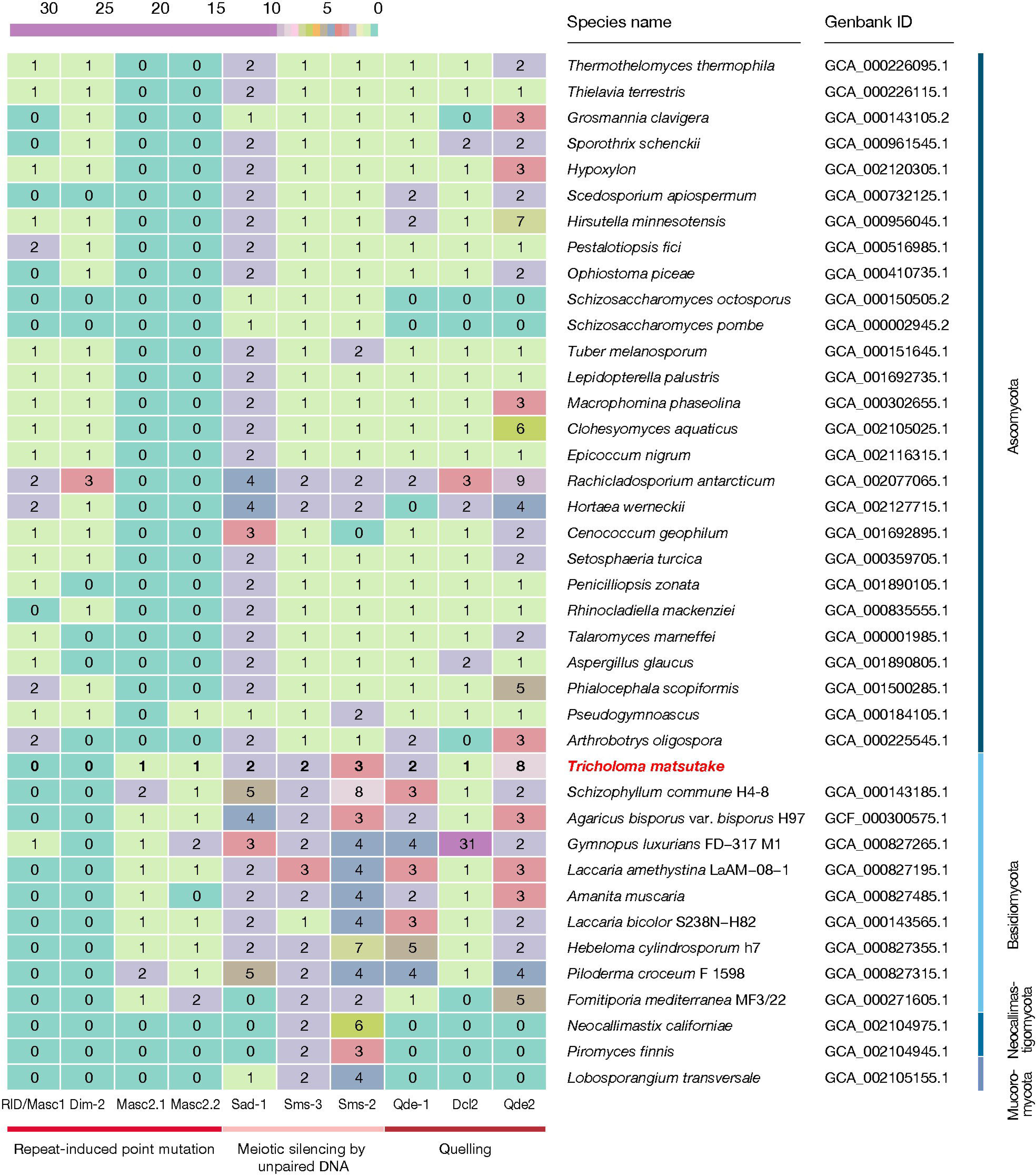
Transposable elements-silencing genes over diverse fungal genomes. The orthologs were inferred by OrthoFinder 1.0.6. Dim-2 and Masc2; Sms-3 and Dcl2; and Sms-2 and Qde2 were further differentiated from gene trees because they belonged to the same gene families. Reference genes are listed in S6 Table.

### Genes enclosed by transposable elements are transcriptionally repressed

It was previously reported that the transcription of TE-surrounded genes is highly repressed in a fungus, *Pleurotus ostreatus* [15]. There were 702 genes surrounded by transposable elements using an ad hoc algorithm described in the Methods section. Among these TE-surrounded genes, the transcripts of 584 genes (83.2%) were never identified at any developmental stage. This was much higher percentage than overall transcriptionally suppressed genes (34.4%). To reveal that these suppressed genes were not pseudogenes or wrongly annotated genes, we investigated their homologs where 152 suppressed genes (26.0%) had paralogous genes that were not enclosed by TEs, and normally expressed in at least one developmental stage (>1 FPKM). In addition, 290 suppressed genes (49.7%) had orthologous genes in at least five Agaricomycetes genome.

### Transcriptomic dynamics in the hyphae, primordia, and fruiting body developmental stages

We compared transcriptomic changes between the hyphae, primordia, and fruiting body developmental stages (Fig 4). Of 15,305 predicted genes, 10,046 (65.6%) genes were transcribed in at least one condition (>1 FPKM) while 5259 genes (34.4%) were never observed in any developmental stage. The majority of the unexpressed genes (4976 genes, 94.6%) were annotated as hypothetical proteins which lacked known functional domains. On the other hand, 355 genes were constantly expressed during development where the genes belonged to top 1000 highest-FPKM genes in all development stages. They were mostly housekeeping genes including ribosomal proteins, heat shock proteins, cytochrome, transporters, and ATP synthases. In the hyphae, primordia, and fruiting body development stages, 2382, 765, and 884 genes were overexpressed over other two stages, respectively (Supplementary Data S1).

**Fig 4.**
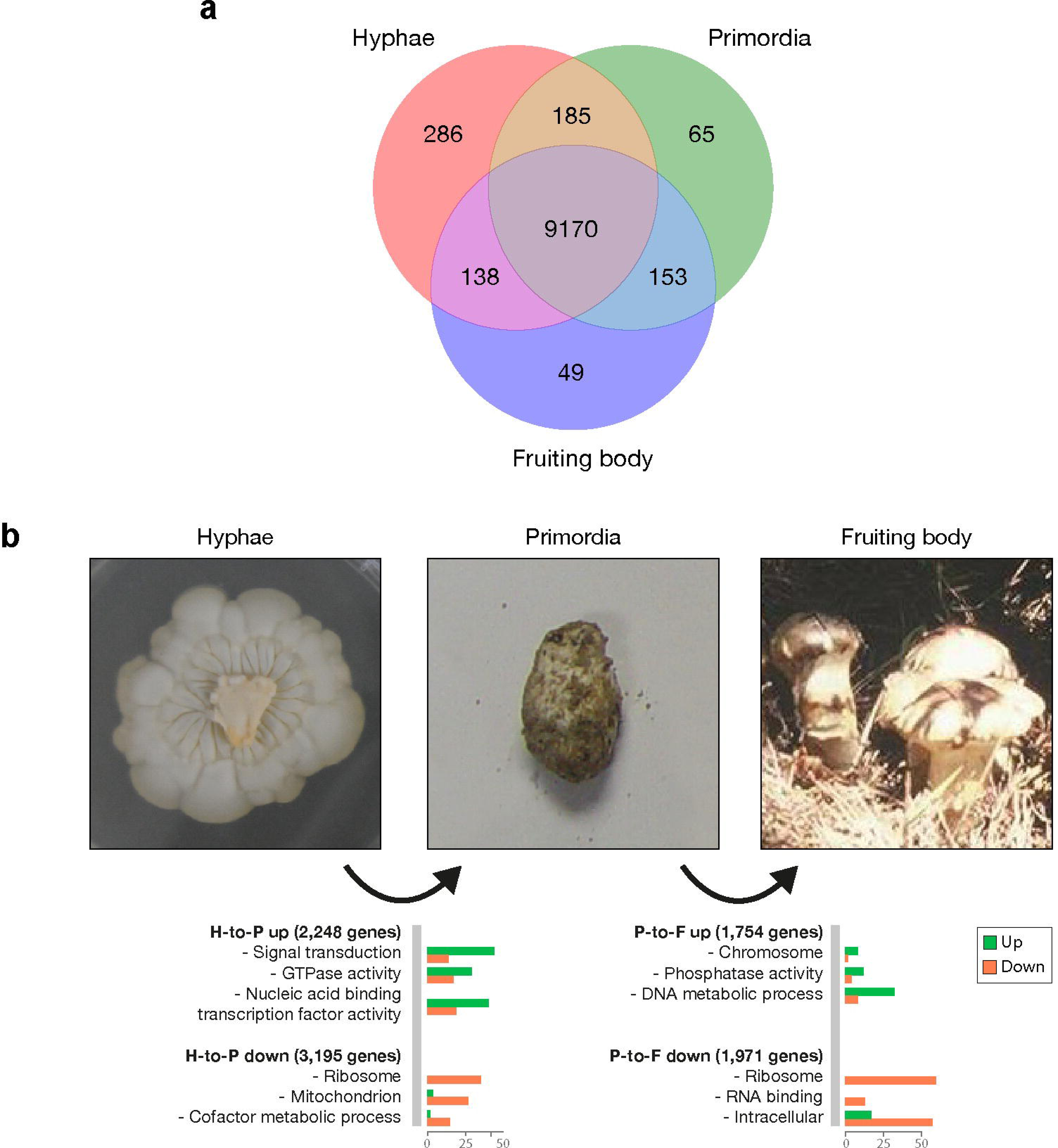
Three developmental stages of *Tricholoma matsutake*: hyphae, primordia, and fruiting body. a. The Venn diagram depicts the number of expressed genes (>1 FPKM) across the three developmental stages. **b.** Up- and down-regulated genes during development. Gene functional categorization was carried out using Gene Ontology Slim (http://www.geneontology.org).

The transition from hyphae to primordia (H-to-P transition) upregulated 2248 genes and downregulated 3195 genes, respectively, while the transition from primordia to fruiting body (P- to-F transition) revealed an upregulation of 1754 genes and a downregulation of 1971 genes (Fig 4). In the H-to-P transition, the gene function related to the signal transduction, GTPase activity, and nucleic acid binding transcription factor activity were enriched, whereas the genes related to ribosome, mitochondrion, and cofactor metabolic process were downregulated. On the other hand, in the P-to-F transition, the enriched functional categories were chromosome, phosphatase activity, and DNA metabolic process, whereas ribosome, RNA binding, and intracellular genes were suppressed (P<0.01 estimated by Fisher’s exact test).

Trima_09940 was the most expressed gene in fruiting body (39299 FPKM). The translated protein had 158 aa length and a signal sequence for secretion. Also, this gene had a diedel domain (PF13164) which is related to insect immune response [22]. The homologs based on sequence similarity were found in *Piloderma croceum* (ectomycorrhizal basidiomycete), *Sistotremastrum niveocremeum* (saprotrophic basidiomycete), *Fusarium mangiferae* (plant pathogenic ascomycete), and so on. The homolog in Drosophila was also identified with 45.8% identity. Although the biological or molecular function of this gene was unclear, it may play essential role in fruiting body formation.

### Transcriptional regulators related to fruiting body formation: transcription factors, light receptors, and hydrophobins

We identified 370 transcription factor genes with DNA-binding domain search. These included *fst4*, *fst3*, *hom1*, *hom2*, *bri1*, *gat1*, and *c2h2* homologs that play critical roles in mushroom formation [7, 23] (S1 Table). While the five genes (*fst4*, *fst3*, *hom1*, *hom2*, and *gat1*) were overexpressed at primordia and fruiting body, as reported in previous studies, the expression levels of *bri1* and *c2h2* were not significantly changed over the three stages. Among differentially expressed 190 transcription factor genes (fold change >2), 53 genes (27.9%) were overexpressed at both primordia and fruiting body (Fig 5). This indicates that these two stages share many regulatory processes over hyphae. The transcription factors overexpressed at primordia and fruiting body were classified as helix-turn-helix, basic helix-loop-helix/leucine zipper, and β-scaffold factors with minor groove contacts.

**Fig 5.**
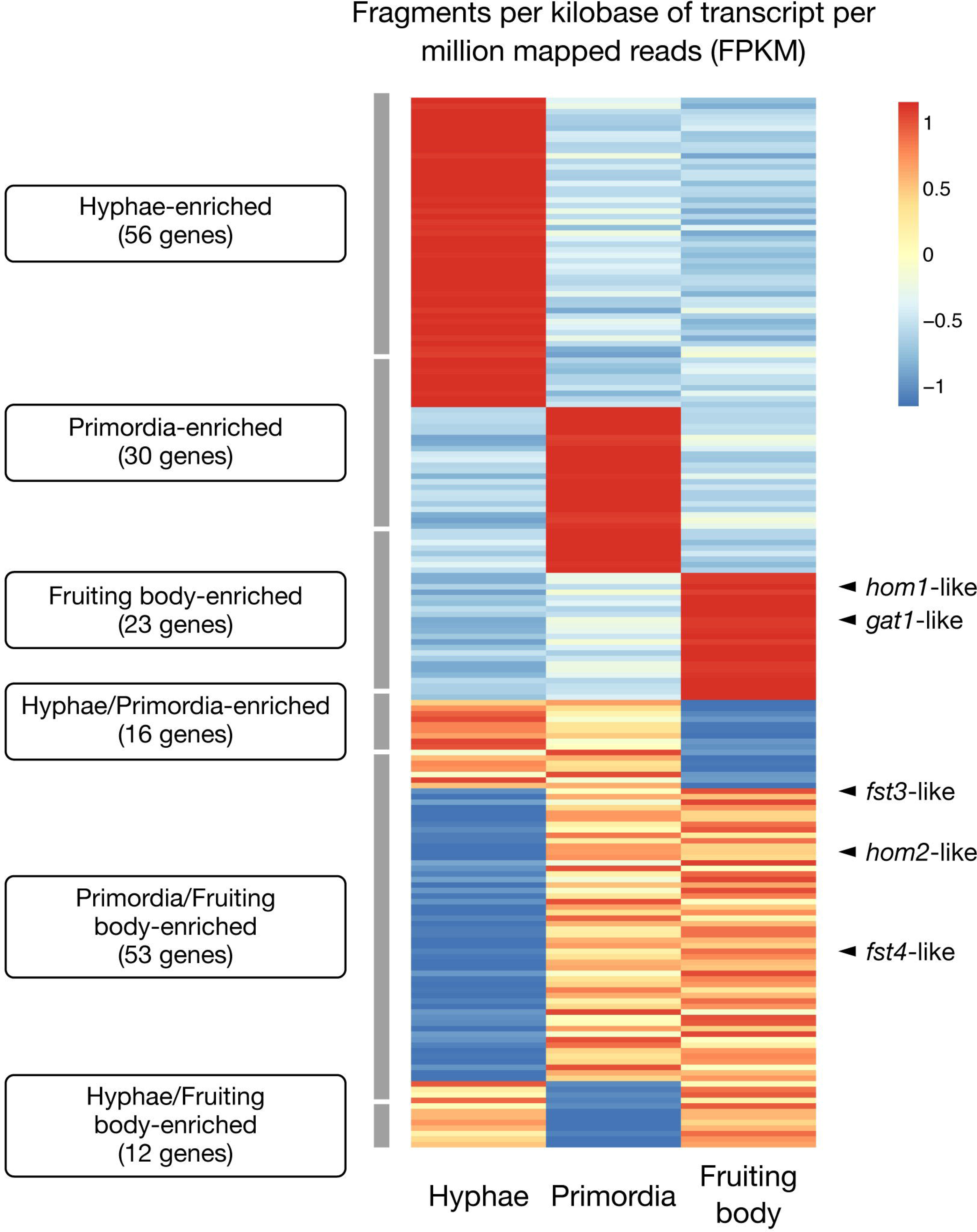
Transcription factor expressions at the three developmental stages. Differentially expressed genes were determined based on logFC (>1 or <-1) calculated by IsoEM2 and IsoDE2. When two conditions were more expressed than the other, but there was no difference between them (-1<logFC<1), we assigned this gene as co-overexpressed in those two conditions. Scaled values based on row Z-scores were used to fill each cell.

Blue light receptor complex WC1/2 is necessary for mushroom development because its deletion prevents mushroom formation [9]. *T. matsutake* harbored the blue light receptor complex WC1/2 (encoded by Trima_13733 and Trima_03536). While *wc1* gene expression was higher in the fruiting body stage, the level of the *wc2* gene was enriched in hyphae and fruiting body (Supplementary Data S1) developmental stage.

Hydrophobins have multiple biological roles that include fruiting body formation and host–fungus interaction [24]. A total of eight hydrophobin genes were annotated, which was relatively small compared with the other 37 Agaricomycete genomes (S6 Fig); it ranged from 0 to 130 (average, 20). All hydrophobin genes were differentially expressed in the three developmental stages: four were only overexpressed at fruiting body, two at hyphae, and one at primordia (S6 Fig). A hydrophobin gene (Trima_02415) was overexpressed at both hyphae and primordia. Trima_15224 had the fifth highest FPKM value in fruiting body among all predicted genes. This hydrophobin gene might be involved in the fruiting body formation.

### Small secreted protein genes are dominantly expressed at hyphae

The small secreted protein genes of ectomycorrhizal fungi are important in the symbiosis development [25]. The number of small secreted protein genes in Agaricomycetes genomes ranged from 196 to 1,053, but the ectomycorrhizal genomes had fewer small secreted protein genes, ranging from 289 to 576 (S7 Fig). Among the 445 predicted small secreted protein genes, 251 genes (56.4%) were differentially expressed with at least one comparison, while 96 (21.6%) genes were not expressed (zero FPKM values) in all stages (S8 Fig). Many of differentially expressed small secreted protein genes were overexpressed at hyphae (87 genes). There were 82 cysteine-rich small secreted protein genes (>3% of cysteine of translated protein sequence), such as fungal specific cysteine rich protein (PF05730), peptidase inhibitor (PF03995), calcium binding protein (PF12192), and various carbohydrate binding modules.

### IstB-like domain is conserved in mycorrhizal genomes

To examine the ectomycorrhizae-specific functional domains, we performed the enrichment test over the *T. matsutake* and 37 Agaricomycetes genomes. Among those 21 Pfam domains enriched in the *T. matsutake* genome (P<0.01 estimated by Fisher’s exact test), IstB-like ATP binding domain (Pfam: PF01695), which is a putative transposase [26], was highly conserved over taxonomically diverse mycorrhizal species (S9 Fig). Although this domain is frequent in bacteria (http://pfam.xfam.org/family/IstB_IS21), fungal IstB-like domains had low sequence similarity with the bacterial domains (<6% matched length coverages). Three were overexpressed in hyphae while one was in fruiting body.

Three additional functional domains were enriched in the *T. matsutake* genome: cadmium resistance transporter (PF03596), Neprosin (PF03080), and carbohydrate binding domain (PF10645, CBM52 family). They were mainly distributed over ascomycete genomes, and only a few basidiomycete genomes possessed them (S10–S12 Figs). Neprosin, a peptidase that cleaves C-terminal to proline residues under highly acidic conditions [27], was usually found in plant genomes, and few bacterial genomes have it. Only the *Tricholoma* and *Laccaria* genomes contained them among fungi. The gene tree of this domain showed the grouping between basidiomycete and several actinobacteria genes (S11 Fig). The CBM52 module is required for septum localization in *Schizosaccharomyces pombe* binding to β-1,3-glucan [28]. This domain was also contained in the *Gymnopus luxurians* and *Sistotremastrum suecicum* genomes with six and one copies, respectively. Further verifications to reveal their exact function remain to be revealed. The functional domains in the *T. matsutake* and the other genomes are summarized in Supplementary Data S2.

### CAZymes are reduced in the genome and differentially expressed during developments

The *T. matsutake* genome had a reduced number of carbohydrate-active enzymes (CAZymes) compared with other Agaricomycetes genomes. A total of 394 predicted CAZyme genes included 143 glycoside hydrolases (GHs), 33 carbohydrate-binding modules (CBMs), 90 glycosyl transferases (GTs), 9 polysaccharide lyases (PLs), 59 carbohydrate esterases (CEs) and 60 auxiliary activities (AAs). The genome had one of the lowest number of total CAZymes compared with other 38 Agaricomycetes. Interestingly, ectomycorrhizal fungi including *T. matsutake* showed similar CAZyme profiles where grouped in a cluster, as seen in Fig 6 except for *Piloderma croceum*. It has been reported that ectomycorrhizal basidiomycetes have lost major gene families, such as plant cell wall-degrading enzymes [13], which includes almost all GH families, especially the GH6 family that enzymatically degrades crystalline cellulose [13, 29]. This lack of the GH6 family was also observed in the *T. matsutake* genome. The CAZymes in the *T. matsutake* and the other genomes are summarized in Supplementary Data S3. Interestingly, we identified two CAZyme submodules uniquely found in *T. matsutake* genome while other ectomycorrhizal genomes lacked: CBM16 and CBM52.

**Fig 6.**
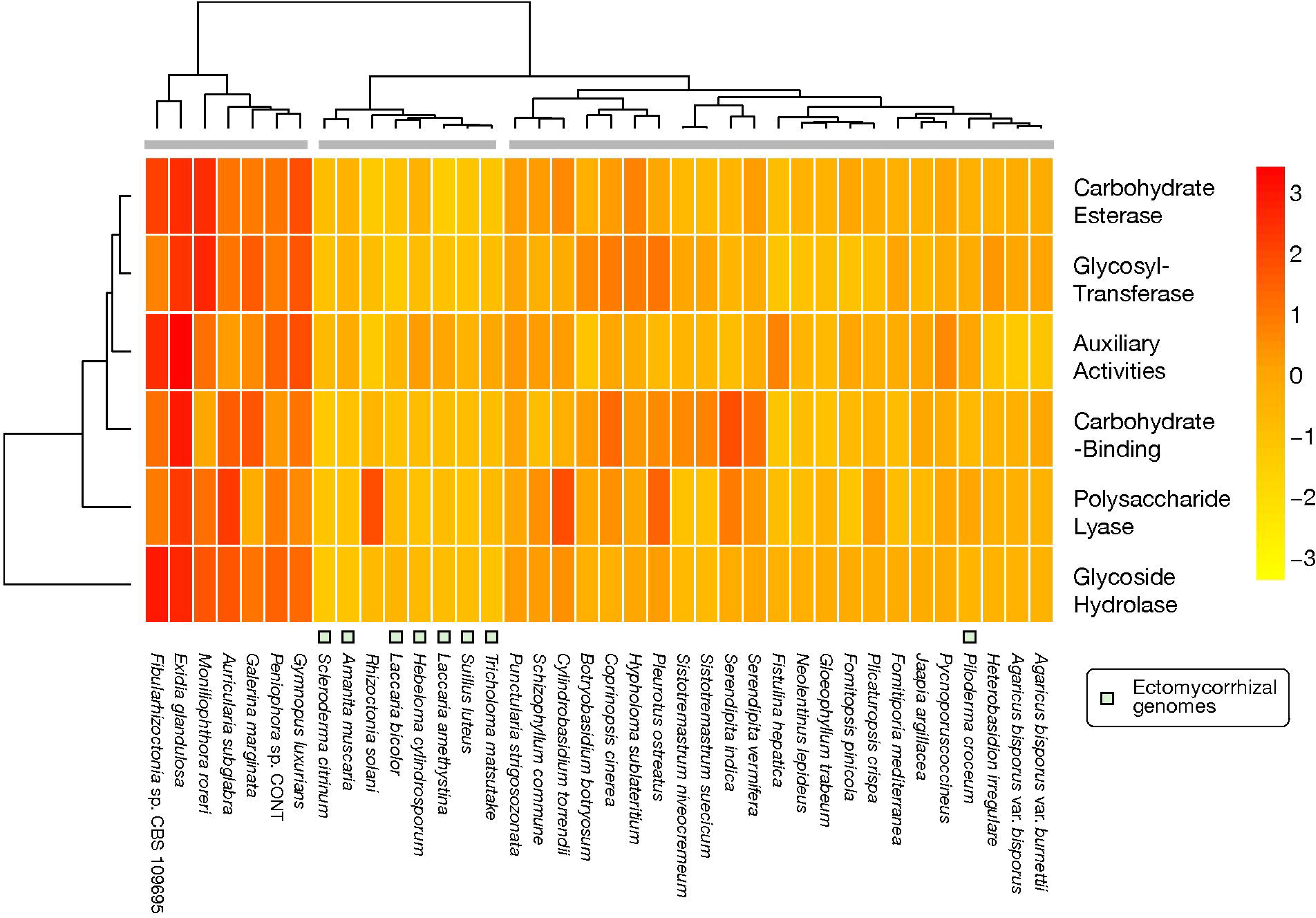
CAZyme genes in the *Tricholoma matsutake* and 37 Agaricomycetes. All available CAZyme modules were counted. Scaled values based on row Z-scores were used to fill each cell.

Different numbers of CAZymes were expressed during development; 57 at hyphae, 44 at primordia, and 47 at the fruiting body were overexpressed (Fig 7). Four of nine PL families were overexpressed in fruiting body. Several GH subfamilies, such as GH5, GH17, and GH20, are only or mostly expressed in fruiting body (4 of 15 for GH5, 2 of 2 for GH17, and 2 of 2 for GH20). GH5 has a role in degradation of lignocellulose [29]. Interestingly, 15 of 44 (34%) primordia-activated genes were auxiliary activities (AAs), while 11 of 57 (19%) and 5 of 47 (11%) were overexpressed at hyphae and fruiting body, respectively. The AA families activated at primordia included AA1, AA3, AA7, and AA9 families.

**Fig 7.**
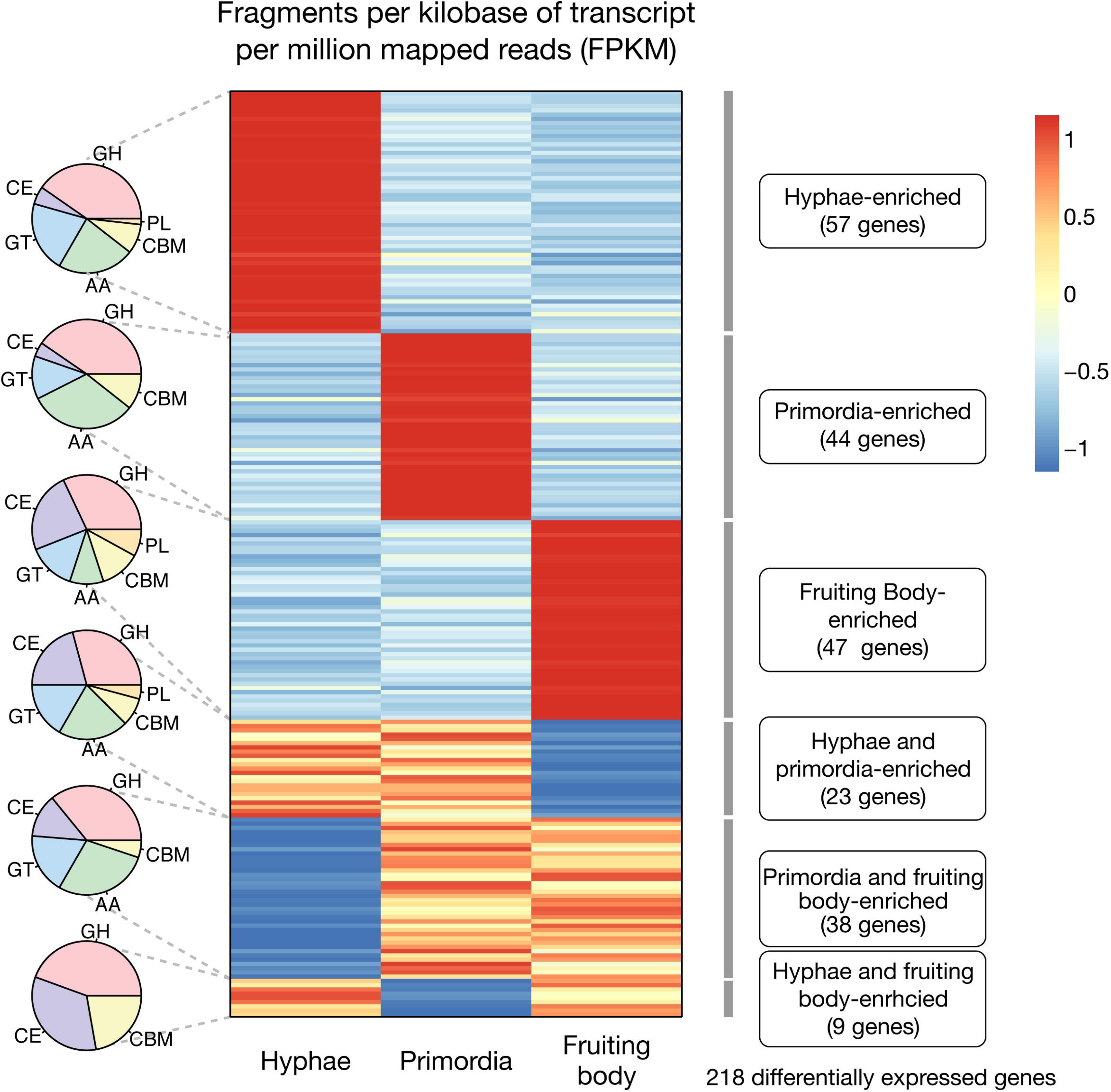
Differentially expressed CAZymes at each of the developmental stages. The pie graphs depict the number of CAZyme families for each specific expression type. Abbreviations for CAZyme families: glycoside hydrolase (GH), carbohydrate-binding module (CBM), glycosyltransferase (GT), polysaccharide lyase (PL), carbohydrate esterase (CE), auxiliary activity (AA). Scaled values based on row Z-scores were used to fill each cell.

## Methods and materials

### Strain and culture conditions

*Tricholoma matsutake* KMCC04578 at the primordia and fruiting body phases were harvested from Gachang, located near Daegu, South Korea. The dikaryotic mycelia were isolated from the gills of the fruiting bodies and cultured in potato dextrose broth (PDB; 4 g/L potato peptone, 20 g/L glucose and pH 5.6 ± 0.2) for 30 days at 25°C.

### Isolation of genomic DNA and total RNA

The genomic DNA was extracted from mycelium, using a cetyl trimethyl ammonium bromide (CTAB)-based fungal DNA isolation protocol [30]. The total RNA was extracted from *T. matsutake* (mycelium, primordium, and stipe of the fruiting body), using an RNA isolation kit (Qiagen, Valencia, CA, USA). The misamples were ground to fine powder with a mortar and pestle under liquid nitrogen. The resulting samples were homogenized with 15 ml of buffer RLT containing β-mercaptoethanol. After centrifugation for 10 min at 3,000 g, the upper phase was mixed with 15 ml of 70% EtOH, and total RNA isolated using an RNA Binding Spin Column under centrifugation for 5 min at 3,000 g. After two wash steps, total RNA was extracted using DEPC-treated water. The RNA samples (A260/A280 >1.8) were collected and subjected to further experiments.

### Genome sequencing and genome assembly

Three sequencing libraries were generated for *T. matsutake*: two Illumina paired-end libraries (500 bp and 250 bp insert size, respectively) and an Illumina mate pair library (5 kbp insert size) (S2 Table). Raw reads were quality-controlled by trimming low quality bases (<30 Phred quality score) and removing short reads after trimming (<50 bp for MiSeq reads and <30 bp for HiSeq reads). Mitochondrial genomic reads were removed by aligning all reads into the reported *T. matsutake* mitogenome sequence (GenBank ID: JX985789.1) [31] using Bowtie2 (*-k 1 --very-sensitive --end-to-end*) [32]. As a result, 1.1% of the total reads were removed.

ALLPATHS [33] was used for the assembly using the three Illumina libraries with *PLOIDY=2* option. We identified and removed four scaffolds derived from vector contamination obtained by BLASTn search against UniVec database (https://www.ncbi.nlm.nih.gov/tools/vecscreen/univec/). One scaffold of human DNA contamination was also removed. Additional sequence contaminations were examined by drawing a scatter plot of GC contents and sequence coverages using Blobology [34] with aligning all scaffolds against NCBI *nt* database. Thus, we corroborated that there was no additional sequence contamination in the final assembly.

### Gene prediction

FunGAP [16] was used to predict protein-coding genes in the assembly. The genome assembly and RNA-seq reads from hyphae were input to the program. *Laccaria_bicolor* gene model was selected for Augustus inside the FunGAP. This generated 17,018 preliminary predicted genes. We manually removed 1,707 transposable element genes such as retrotransposon gag protein (Pfam: PF03732) and reverse transcriptase (Pfam: PF07727) based on Pfam annotation by InterProScan 5.25-64 [35]. Targeted Pfam domains are listed in S3 Table.

The remaining 15,305 genes were examined for their reliability by RNA-seq reads alignment, functional domain annotation, and ortholog search against relatives. RNA-seq reads from three development stages (hyphae, primordia, and fruiting body) were aligned into the genome assembly using HISAT2 [36] and FPKM values were calculated using IsoEM2 [37]. Only >1 FPKM genes were considered as RNA-seq-supported genes. Pfam domains were annotated by InterProScan 5.25-64 and the genes containing at least one Pfam domain were considered as functional domain-supported genes. Ortholog search was performed by OrthoFinder 1.0.6 [38] using *T. matsutake* and 37 Agaricomycetes genomes. When a gene belongs a gene family that contains members from more than five genomes, we considered that the gene is supported by ortholog search. To check for genome completeness, we used BUSCO v3.0.2 [17], in which the *basidiomycota_odb9* database was used. Because the assembled genome was dikaryotic, we estimated how many genes are allelic by comparing the numbers of two-member gene families with five Agaricales genome. This was obtained by parsing OrthoFinder output.

### Comparative analysis

We chose 37 Agaricomycetes genomes for comparative analyses (S4 Table). In the NCBI database, there are 59 Agaricomycetes genome assemblies with predicted genes as of time of writing this article. We eliminated three incomplete genomes based on BUSCO calculations (<90% completeness). We also sampled two genomes from each order, excluding Agaricales (which *T. matsutake* belongs to) to reduce computing time. This yielded 38 genomes as the final targets for comparative analysis. A species tree was built using RAxML 8.1.3 [39] from concatenated single-copy orthologs obtained by OrthoFinder 1.0.6 [38]. We used *-f a -x 12345 -p 12345 -# 100 -m PROTGAMMAWAG* options for RAxML. Mafft 7.273 [40] and Gblocks 0.91b were used to align the concatenated sequences and extract the conserved regions.

### RNA-sequencing

Illumina RNA sequencing generated 108, 133, and 127 million RNA-seq reads from hyphae, primordia, and fruiting body, respectively (S5 Table). Trim Galore 0.4.4 (https://www.bioinformatics.babraham.ac.uk/projects/trim_galore/) was used for adapter removal, low-quality base trimming (<20 phred score), and short reads filtering (<40 bp). The RNA-seq reads were aligned to the genome by HISAT 2.0.5 [36]. Due to a single replicate of RNA-sequencing libraries, differentially expressed genes were estimated by IsoEM2 and IsoDE2, performing bootstrapping-based approach using an accurate expectation-maximization algorithm [37]. We used *--auto-fragment-distrib* option for IsoEM2 and *-pval 0.05* (desired P value) option for IsoDE2. These programs generate a logarithm of fold change (logFC) between two conditions for each gene. When the logFC value of certain gene is less than -1 or greater than 1, we considered this gene is significantly differentially expressed. We classified all the genes into the seven expression patterns: hyphae-enriched, primordia-enriched, fruiting body-enriched, hyphae and primordia-enriched, primordia and fruiting body-enriched, hyphae and fruiting-body-enriched, and not significantly different among the samples. When two conditions were more expressed than the other, but there was no difference between them, we assigned this gene as co-overexpressed in those two conditions.

To obtain upregulated or downregulated genes during development, we accounted for differentially expressed genes in the hyphae-primordia comparison and the primordia-fruiting body comparison. The differentially expressed genes were functionally classified based on Gene Ontology terms. First, Gene Ontology terms were assigned by running InterProScan 5.25-64 [35] against PfamA database with *--goterms* option. Second, we assigned each Gene Ontology term to high level Gene Ontology term (GO slim) by running owltools (http://code.google.com/p/owltools/) with *--map2slim* and *--subset goslim_generic* options. Finally, Fisher’s exact enrichment test on each GO slim using Python *scipy.stats.fisher_exact* function (https://www.scipy.org/).

### Transcription factor annotation

Transcription factor genes were predicted based on the Pfam domain annotation. Transcription factor Pfam domains were obtained from DBD database (http://www.transcriptionfactor.org) [41] in addition to three functional domains: ARID/BRIGHT DNA binding domain (PF01388) and fungal specific transcription factor domains (PF04082 and PF11951). Known transcription factor homologs were identified by BLASTp search where the best hit was selected. Their orthologies were validated using OrthoFinder 1.0.6 [38].

### Functional domain annotation

InterProScan 5.25-64 [35] predicted functional domains from the protein sequences of *T. matsutake* and 38 Agaricomycetes genomes with Pfam 31.0 [43]. The enriched and depleted functions were estimated by Fisher’s exact test with *scipy.stats.fisher_exact* function of Scipy Python module (https://www.scipy.org/). Four selected functions (PF01695, PF03596, PF03080, and PF10645) were BLASTp-searched against NCBI *nr* database and 50 top hit sequences were used to build the gene trees. Mafft 7.273 [40] was used for multiple genome sequence alignment with *--maxiterate 1000 --localpair* options. FastTree 2.1.10 [44] built the trees with default options.

### CAZyme annotation

Carbohydrate-related enzymes were predicted by three different tools: dbCAN HMMs 5.0 [45], a database that uses HMM profiles of known CAZyme sequences; BLASTp, a tool for searching the protein sequences against the CAZyme sequence database; and Pfam 31.0, domains annotated with CAZyme entries. All three tools were run and integrated to make a final CAZyme prediction. We assigned a CAZyme when more than two of the programs gave the same prediction on a gene.

### Small secreted protein gene prediction

Referring to previous works [46–48], we combined four extracellular proteins prediction programs: SignalP, WoLF PSORT, TargetP, and ProtComp. SignalP 4.1 [49] was run with default option and “signal peptide=Y” and “Networks-used=SignalP-noTM” tags were used to get signal peptide-containing protein sequences. WoLF PSORT [50] was used with “OrganismType=fungi” and most voted localization was used for each protein. TargetP 1.1 [51] was used with -N option for using non-plant networks and “Loc=S” was used to get secreted proteins. ProtComp v9 (http://www.softberry.com/berry.phtml) was used with *-NODB -NOOL* options and “Integral Prediction of protein location” was used for assigning protein locations.

Four programs predicted 780, 1344, 2173, and 823 proteins as secreted, respectively (S7 Fig). Only 78 proteins were predicted as secreted by all programs. The genes predicted by at least three programs (589 proteins) were considered as preliminary secreted protein. We excluded transmembrane, endoplasmic reticulum, glycophosphatidylinositol-anchored proteins from the candidates with TMHMM, PS_SCAN, and GPI-SOM programs. TMHMM 2.0 [52] was used with default options, and when a transmembrane helix was located at within 70 aa N-terminal and other helixes not identified, we considered this as a non-transmembrane protein as referring to previous work [53]. PS_SCAN 1.86 [54] was used to scan endoplasmic reticulum targeting-proteins (PROSITE: PS00014). GPI-SOM 1.5 [55] was used with default options. This resulted in 1788 transmembrane proteins, 16 endoplasmic reticulum-targeting proteins, and 1718 glycophosphatidylinositol-anchored proteins. From the removing of these proteins, we obtained 455 proteins as secreted protein in the *T. matsutake* genome.

### Repeat elements analysis

RepeatModeler and RepeatMasker (http://www.repeatmasker.org) were sequentially used to predict repeat elements in the genomes. Protein-coding sequences within repeat elements were predicted by running Braker1 [56] on the unmasked assembly. The evidence of repeat-induced point mutations was calculated by following the Amselem et al.’s (2015) method [21]. Briefly, repeat sequences were extracted into a FASTA file using *rmOut2Fasta.pl* script within the RepeatMasker package. We split the sequences so that one FASTA file would contain one repeat family sequences. C-to-T hypermutation of specific dinucleotides was calculated using Mafft-7.273 [40] and RIPCAL 2.0 [57]. Finally, the dinucleotide biases were calculated by counting repeat elements with >2 transition/transversion ratio and when more than one-third of the sequences had dinucleotide hypermutations bias. We considered TE-surrounded genes when a gene has repeat elements at both upstream and downstream within 1000 bp distance. Only >400 bp repeat elements were accounted for because there were so many short fragments.

The genes responsible for genome defense against transposable elements were identified using gene family and gene tree analyses. We targeted three mechanisms, including repeat-induced point mutation, meiotic silencing by unpaired DNA, and quelling; nine reference genes related to these mechanisms were used to find their orthologs in the proteome of *T. matsutake* and the other genomes (S6 Table). We identified gene families of BLASTp top hits against the reference genes. The genes of Dim-2 and Masc2; Sms-3 and Dcl2; and Sms-2 and Qde2 belonged to the same gene family; therefore, we additionally constructed gene trees to distinguish them. Mafft-7.273 [40] and FastTree [44] were used to build the gene trees.

## Conclusion

This study aims to understand the genomic circumstances of ectomycorrhizal *Tricholoma matsutake*. The species had the unusually large genome size from repetitive transposable elements insertions. These inserted transposable elements occasionally are involved in transcriptional suppression of nearby protein-encoding genes. We identified the evidence of genome defense against transposable elements by C to T hypermutation with a bias over ’CpG’ dinucleotides. Developmental transcriptomic dynamics revealed that many transcriptional factors are expressed in primordia and fruiting body while small secreted proteins in hyphae stage. The genome contained less carbohydrate-active enzymes as other ectomycorrhizal fungi. These results will help understand how *Tricholoma matsutake* has developed and maintained its lifestyle.

## Data availability

This Whole Genome Shotgun project has been deposited at DDBJ/ENA/GenBank under the accession PKSN00000000. The version described in this paper is version PKSN02000000. MycoBank ID of this species is 307044. Genome browser and data download are available at http://compbio.korea.ac.kr/tricholoma. The authors declare that all other data supporting the findings of this study are available within the article and its Supplementary Information files or are available from the corresponding authors upon request.

---

**S1 Fig.**
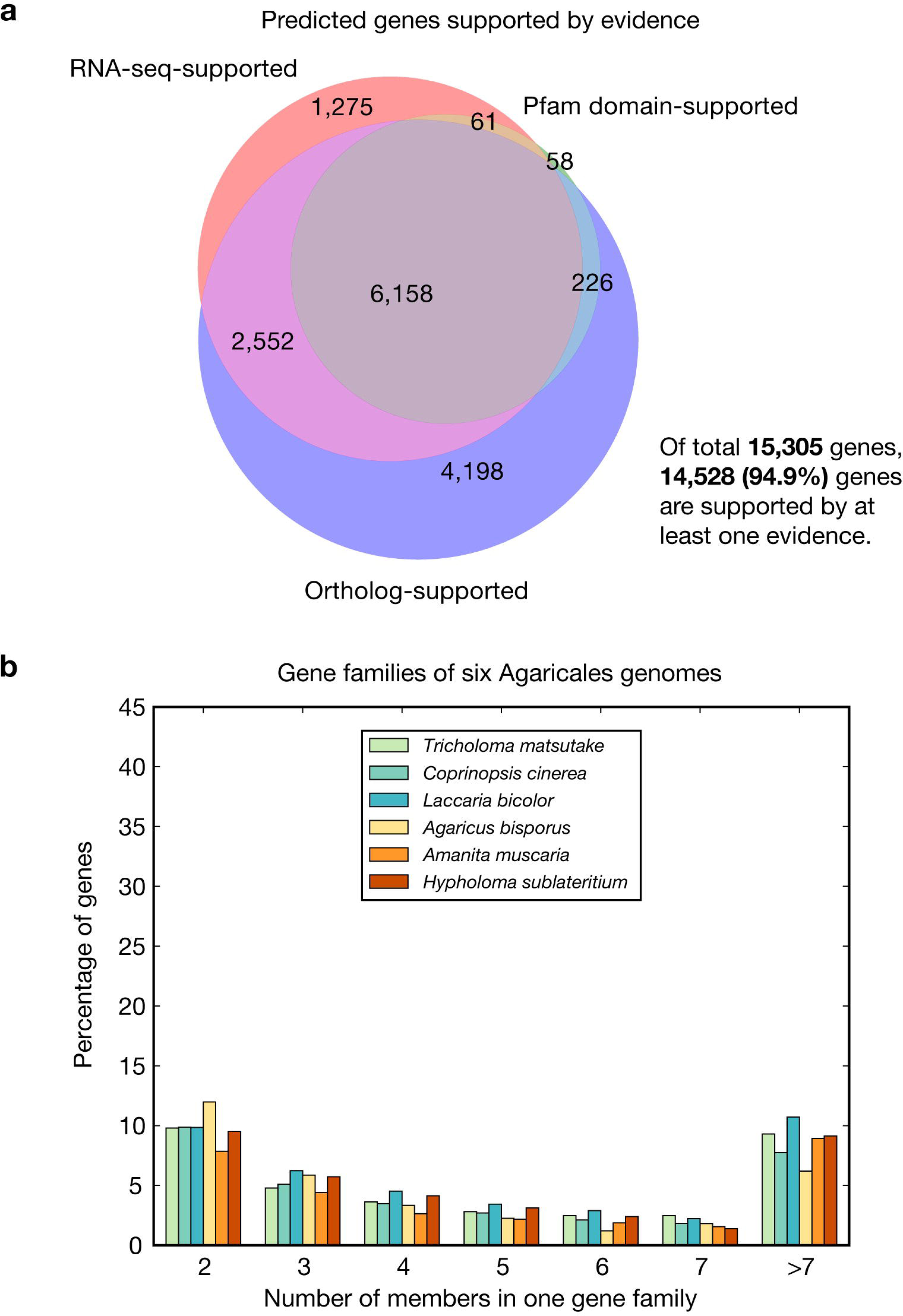
Gene prediction quality assessment and allele genes estimation. a. The predicted genes were examined for their reliability using RNA-seq, Pfam domains, and orthologs within 37 Agaricomycetes genomes. For RNA-seq, we accounted for the genes with >1 FPKM in at least one development stage. When a gene family includes members from more than five genomes, we considered those members are reliable (not pseudogene). **b.** The frequencies of gene members in a gene family.

**S2 Fig.**
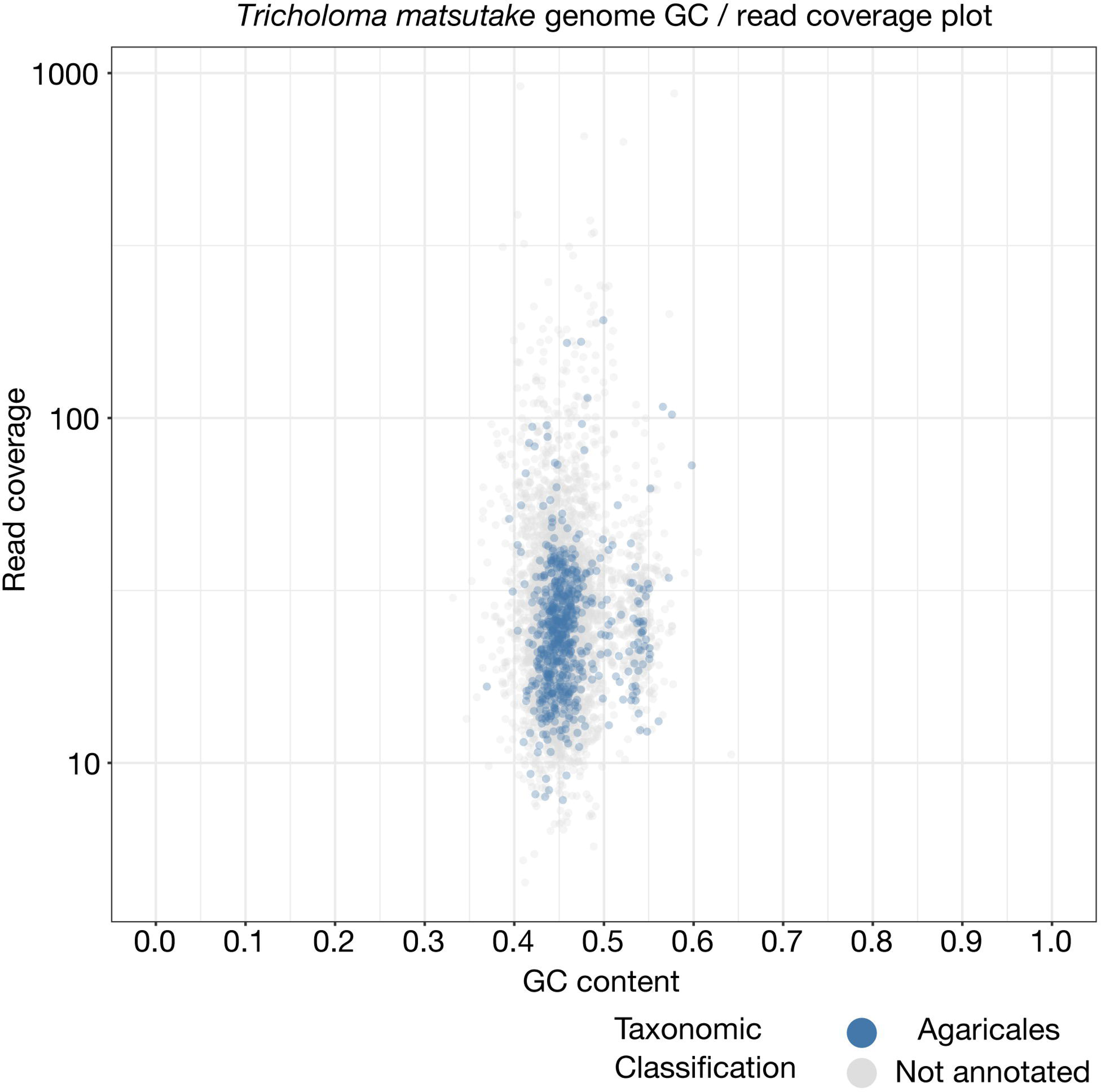
A scatter plot of GC and read coverage of the *Tricholoma matsutake* genome. Each point indicates a scaffold marked with a color according to its taxonomic assignment (order level) which was made based on BLASTn search against the NCBI *nt* database.

**S3 Fig.**
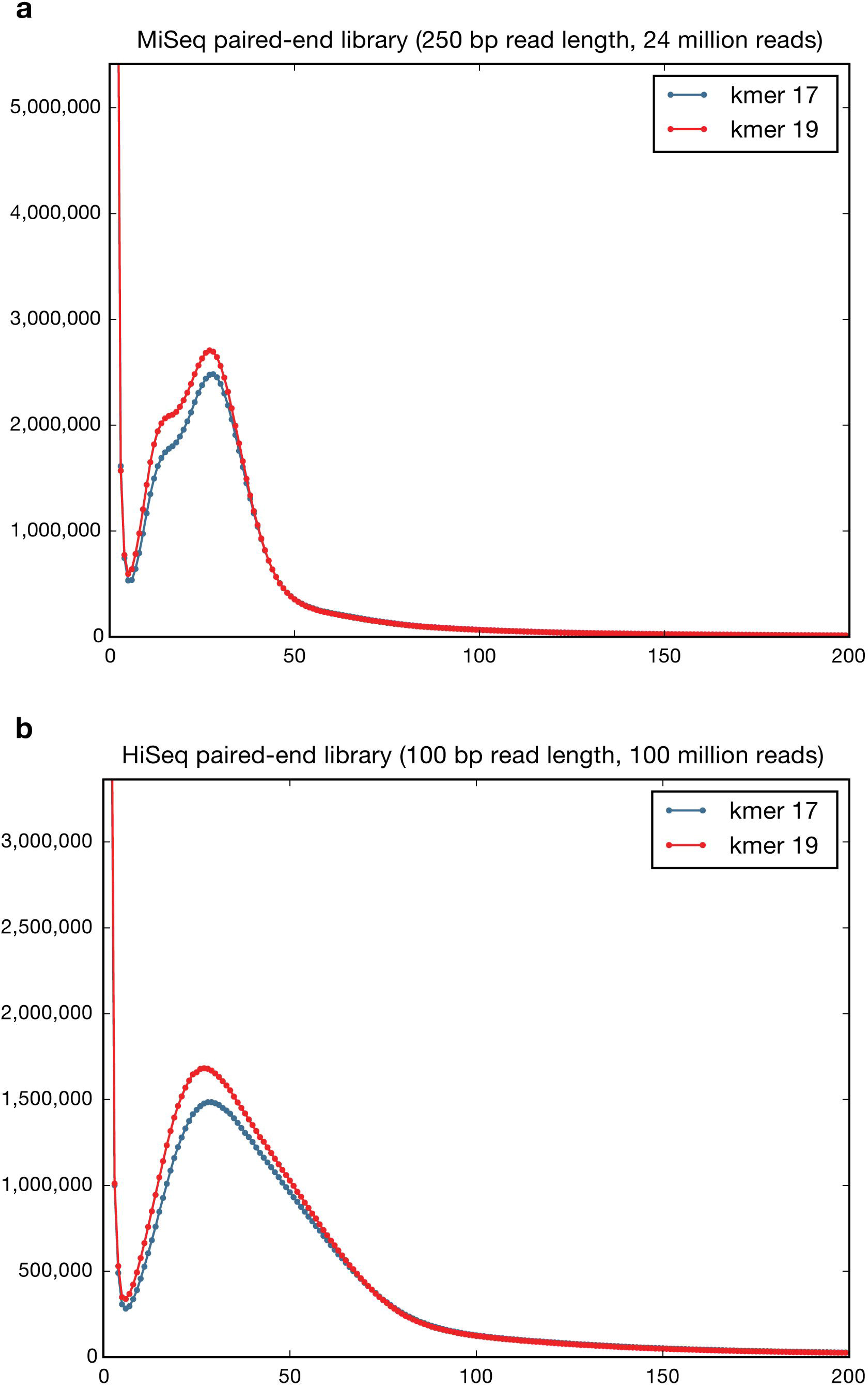
K-mer frequency of genomic DNA sequencing. Two paired-end sequencing libraries were used.

**S4 Fig.**
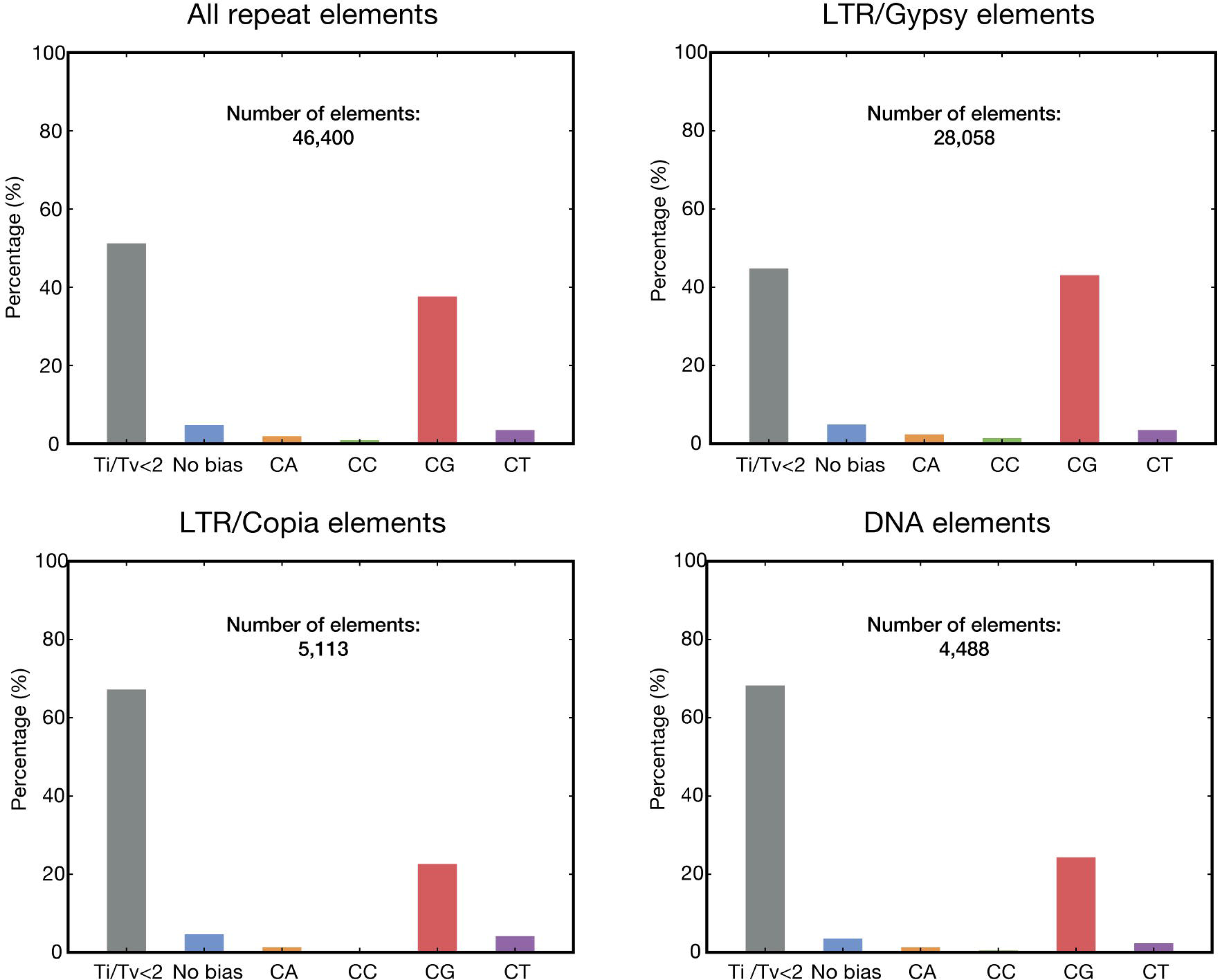
The evidence of repeat-induced point mutation in the *T. matsutake* genome. The repeat elements showing transition/transversion ratio >2 were classified according to the dinucleotide hypermutation bias to see which dinucleotide is preferred in C to T transition during RIP process.

**S5 Fig.**
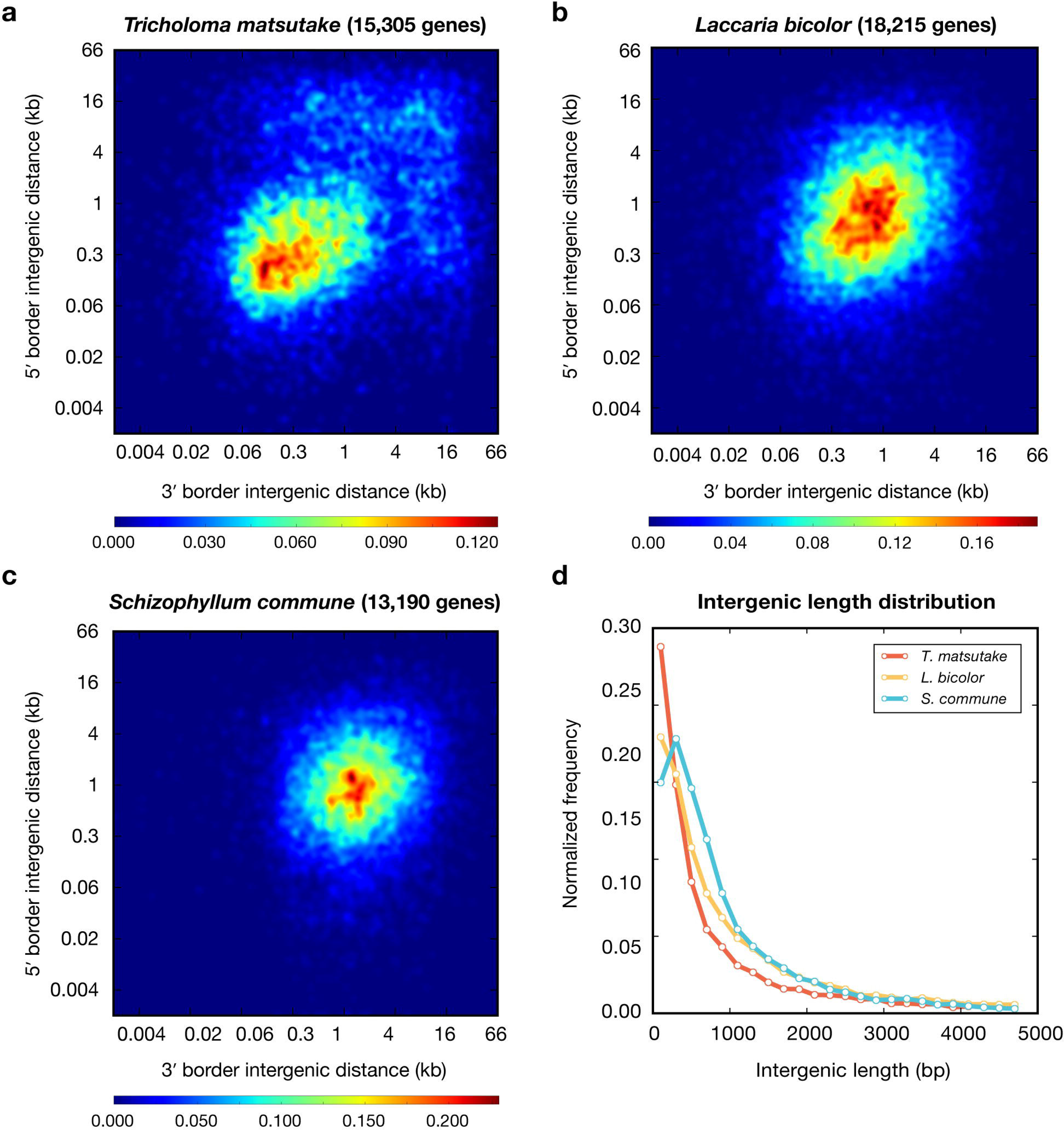
Intergenic length distribution. a–c. GFF3 files were used to obtain the distances to neighbor genes. Both ends of genes of scaffolds were not considered because they had only one side border. Two-dimensional histograms were calculated with 1000 bins and Gaussian filter was used to smooth the heatmaps. **d.** Intergenic length distribution of three mushroom genomes. Frequency values were normalized by total gene counts of the genomes.

**S6 Fig.**
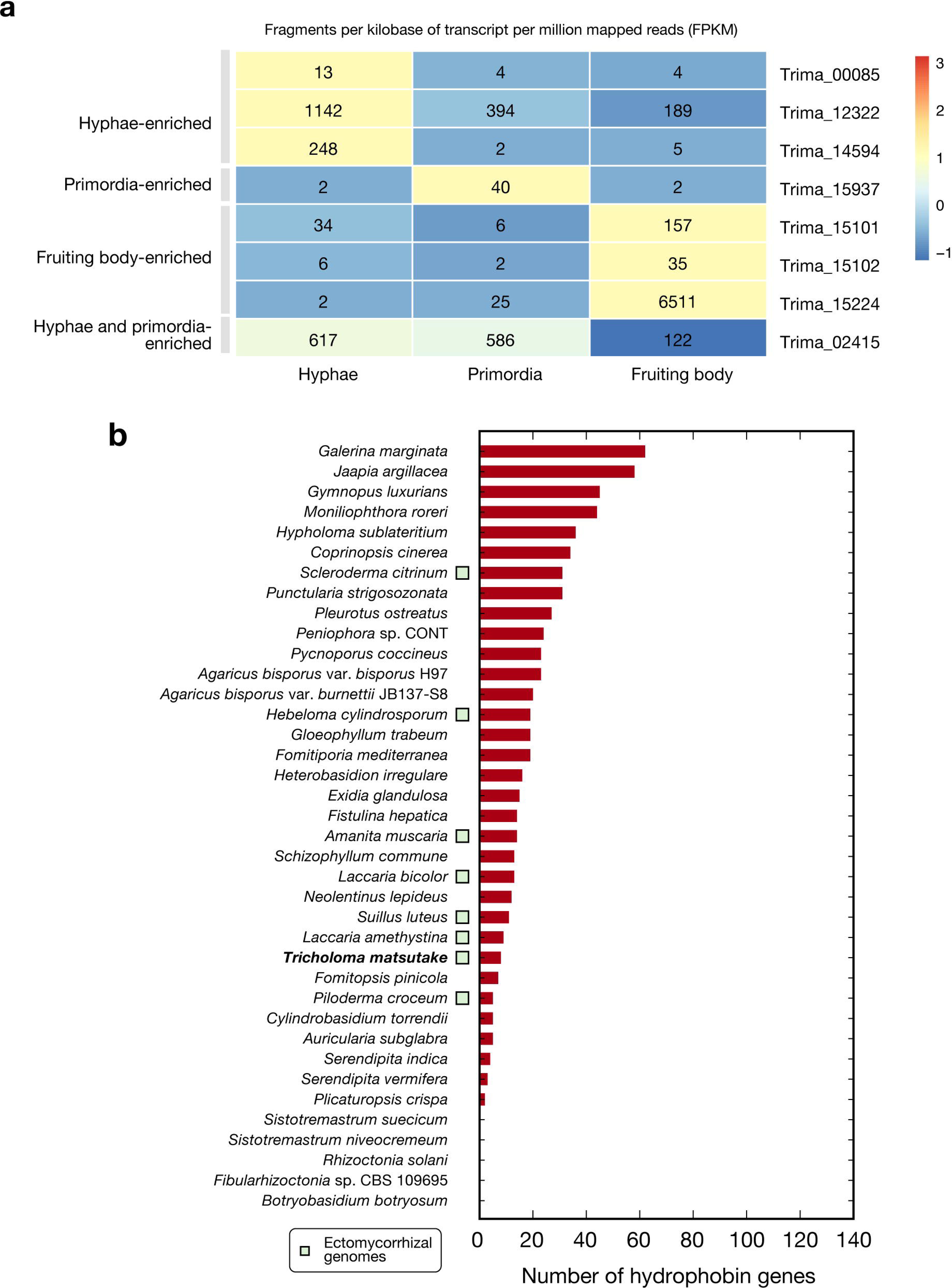
Hydrophobin gene expression at the three developmental stages and the gene frequencies of the 37 Agaricomycetes genomes. Differentially expressed gene types were determined based on logFC (>1 or <-1) estimated by IsoDE2, based on bootstrapping of aligned reads. When two conditions were more expressed than the other, but there was no difference between them (-1<logFC<1), we assigned this gene as co-overexpressed in those two conditions. Scaled values based on row Z-scores were used to fill each cell. The genomes list is shown in Supplementary Table S3.

**S7 Fig.**
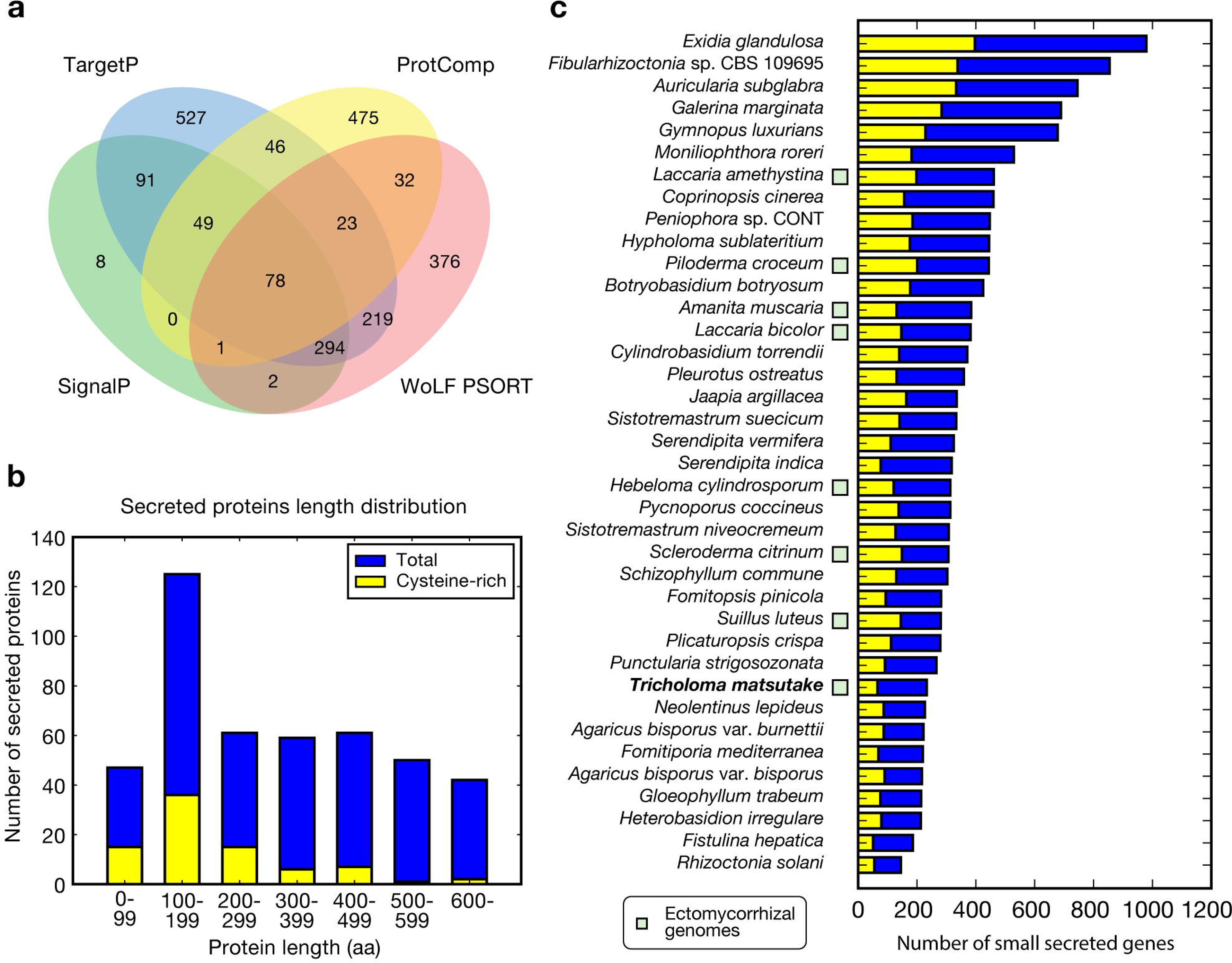
Secreted protein genes. a. The Venn diagram of predicted secreted protein genes predicted by four different programs. **b.** Length distribution of secreted proteins of the *T. matsutake* genome. Cysteine-rich protein sequences are determined when they contain >3% cysteines in the sequence. **c.** Secreted small protein genes (<300 aa) of 38 Agaricomycetes genomes.

**S8 Fig.**
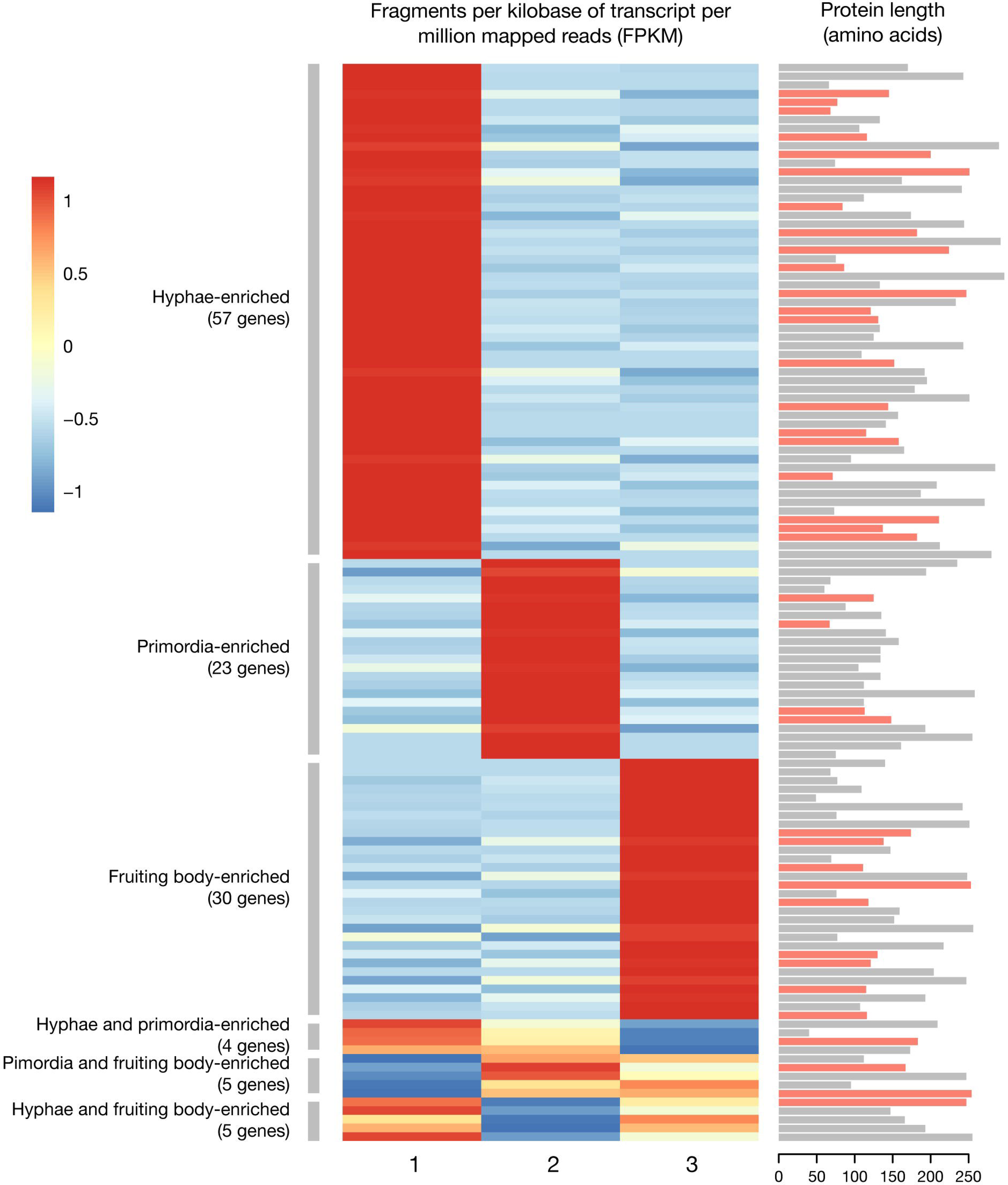
Differentially expressed small secreted protein genes. Differentially expressed genes were determined by IsoDE2, based on bootstrapping of aligned reads. Less than 300 aa length of proteins were considered. Cysteine-rich proteins (>3% of a protein sequence) were marked with light pink color at the protein length bars. Scaled values based on row Z-scores were used to fill each cell.

**S9 Fig.**
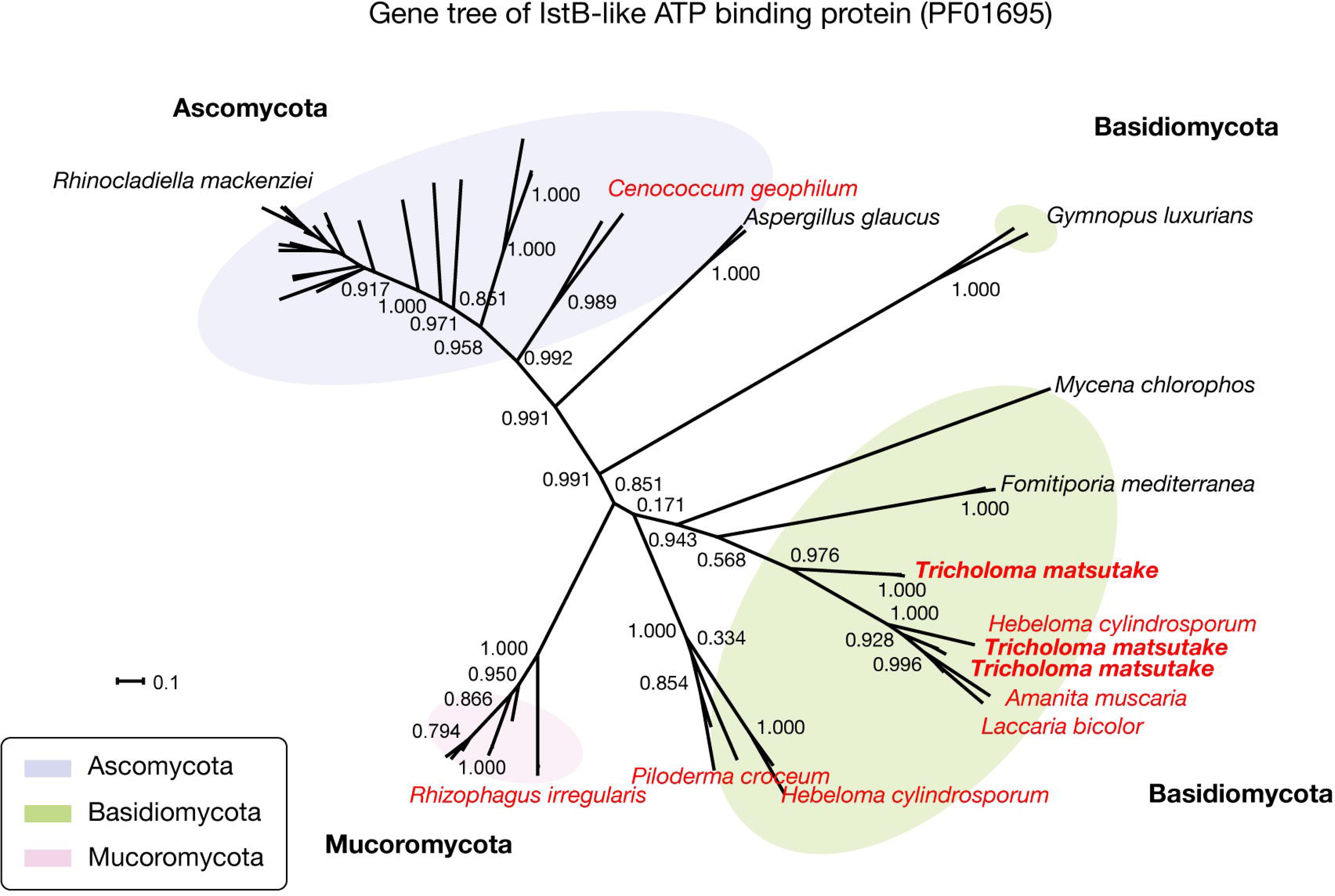
Gene tree of IstB-like ATP binding protein. The 50 homologs were retrieved from BLAST search against the NCBI *nr* database. The gene tree was built using Mafft 7.273 for multiple sequence alignment and FastTree for tree building. The scale bar that represents the mean number of amino acid substitutions per site is shown. Local support values computed with the Shimodaira-Hasegawa test are indicated. Mycorrhizal species are marked in red color.

**S10 Fig.**
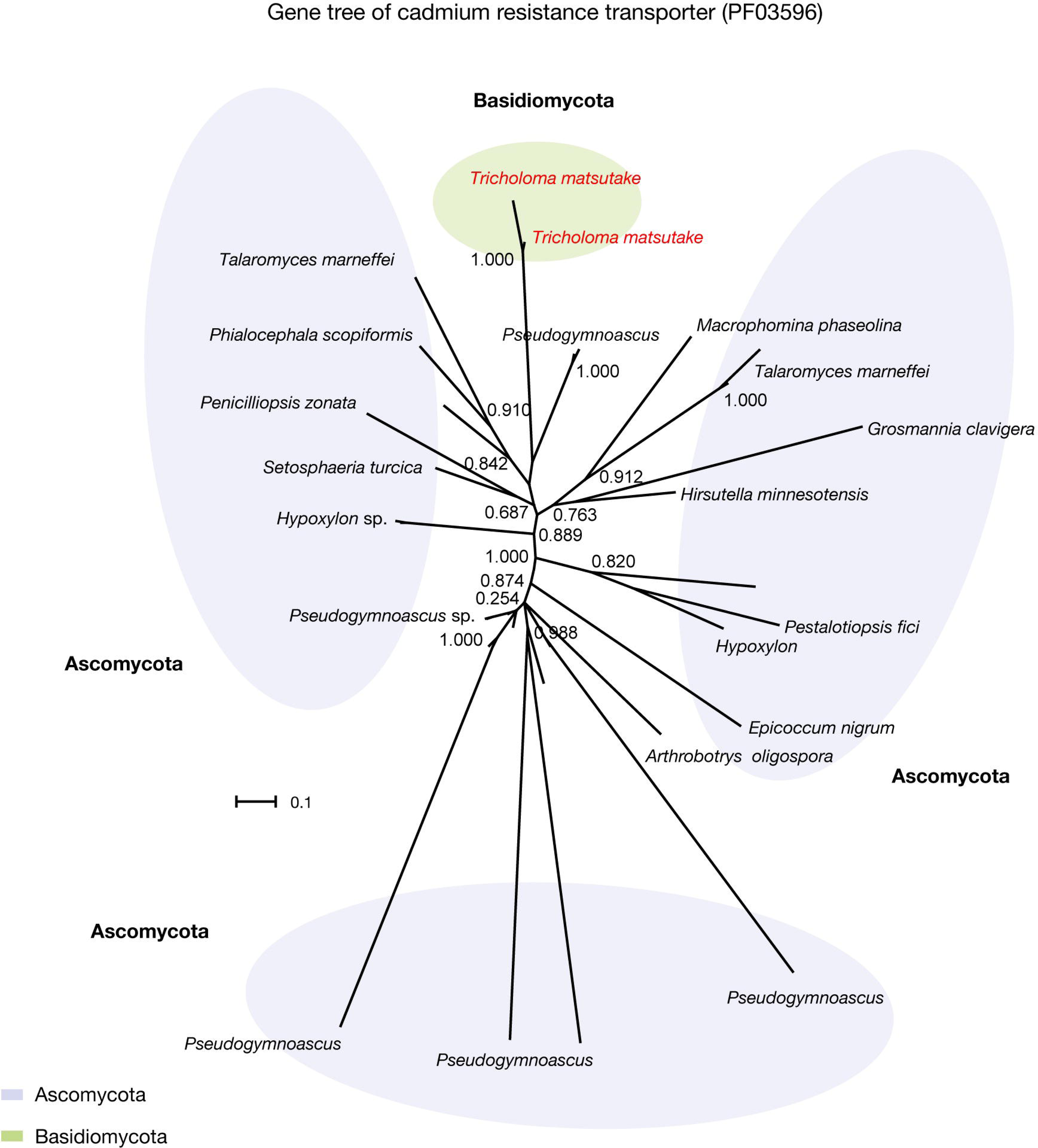
Gene tree of cadmium resistance transporter. The 50 homologs were retrieved from BLAST search against the NCBI *nr* database. The gene tree was built using Mafft 7.273 for multiple sequence alignment and FastTree for tree building. Local support values computed with the Shimodaira-Hasegawa test are indicated. The scale bar that represents the mean number of amino acid substitutions per site is shown. *Tricholoma matsutake* is marked with red color.

**S11 Fig.**
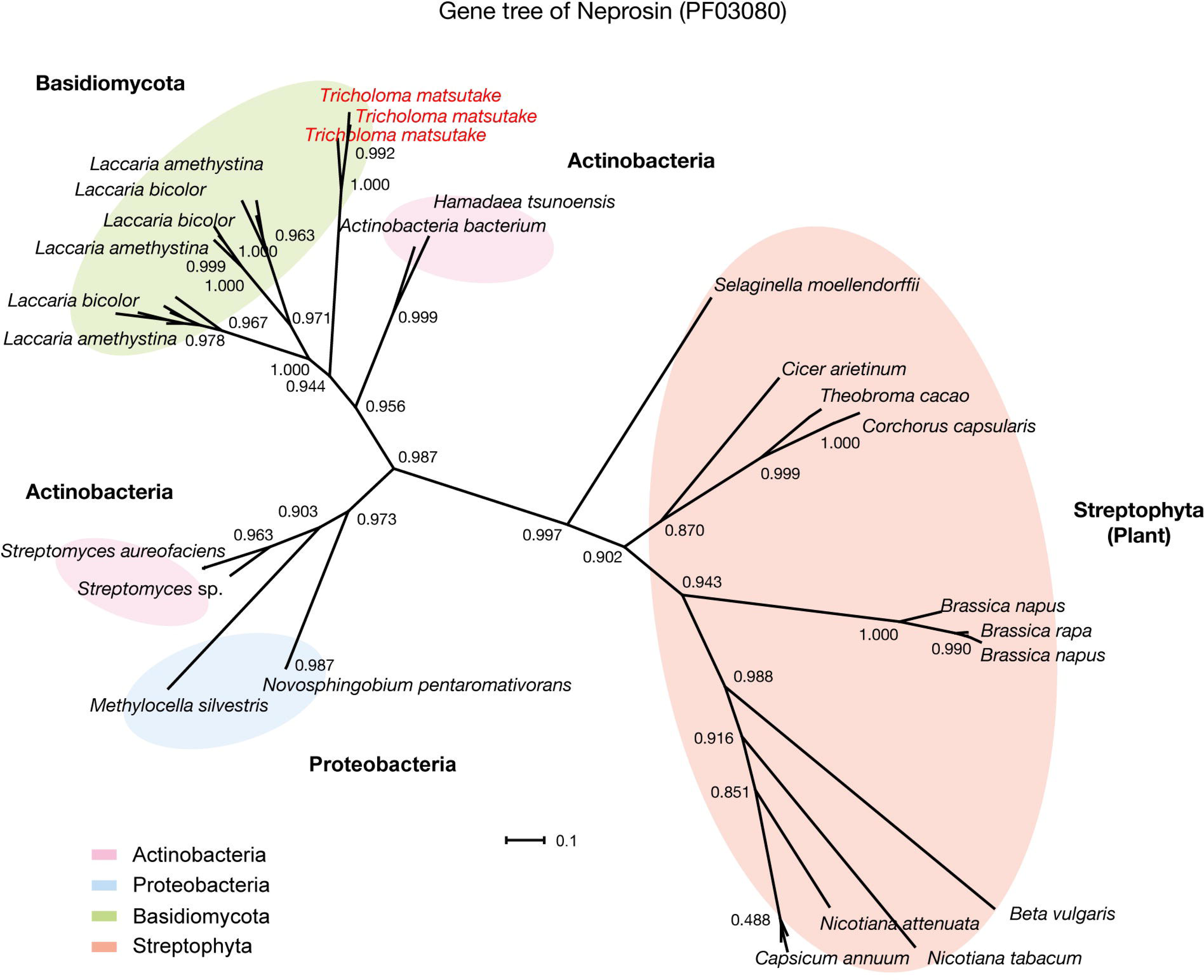
Gene tree of neprosin. The 50 homologs were retrieved from BLAST search against the NCBI *nr* database. The gene tree was built using Mafft 7.273 for multiple sequence alignment and FastTree for tree building. Local support values computed with the Shimodaira-Hasegawa test are indicated for each branch. The scale bar that represents the mean number of amino acid substitutions per site is shown. *Tricholoma matsutake* is marked with red color.

**S12 Fig.**
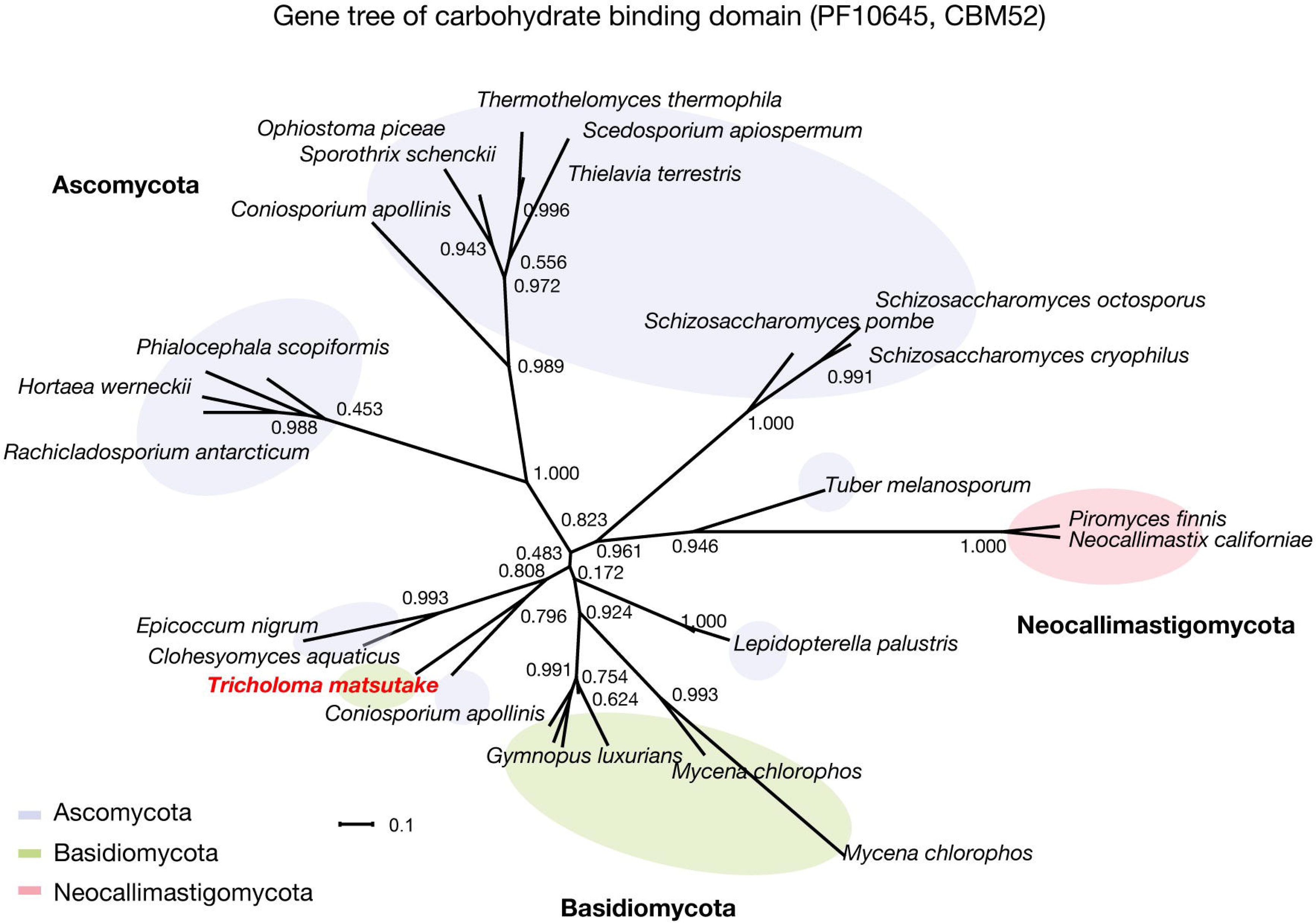
Gene tree of carbohydrate binding domain. The 30 homologs were retrieved from BLAST search against the NCBI *nr* database. The gene tree was built using Mafft 7.273 for multiple sequence alignment and FastTree for tree building. Local support values computed with the Shimodaira-Hasegawa test are indicated for each branch. The scale bar that represents the mean number of amino acid substitutions per site is shown. *Tricholoma matsutake* is marked with red color.

S1 Table. Transcription factors involved in mushroom formation

S2 Table. The summary of genome sequencing

S3 Table. Transposable element Pfam domains

S4 Table. Genomes used for comparative analysis

S5 Table. Total bases and read counts for the three developmental stages of *Tricholoma matsutake*

S6 Table. Target proteins responsible for genome defense by fungi

S1 Data. mRNA expressions.

S2 Data. Functional domains of *T. matsutake* and Agaricomycetes genomes.

S3 Data. CAZymes of *T. matsutake* and Agaricomycetes genomes.

