## Supplementary material for "Unusual genome expansion and transcription suppression in ectomycorrhizal *Tricholoma matsutake* by repetitive insertions of transposable elements": S1 Table

**S1 Table. Transcription factors involved in mushroom formation**

| Protein name | GenBank Access | Length  (aa) | Homologs in *Tricholoma matsutake* | Length  (aa) | E-value | Identity  (%) |
| --- | --- | --- | --- | --- | --- | --- |
| Hom2 | XP_003029756.1 | 723 | Trima_14430 | 679 | 2.49e-57 | 33.4 |
| Bri1 | XP_003038897.1 | 776 | Trima_08826 | 932 | 1.99e-159 | 39.2 |
| Fst4 | XP_003034563.1 | 826 | Trima_15097 | 859 | 0.0 | 45.1 |
| C2H2 | XP_003026630.1 | 360 | Trima_03635 | 486 | 7.59e-32 | 61.9 |
| Fst3 | XP_003031320.1 | 1120 | Trima_16495 | 1056 | 0.0 | 64.8 |
| Gat1 | XP_003036589.1 | 449 | Trima_00186 | 527 | 6.07e-66 | 39.1 |
| Hom1 | XP_003030056.1 | 316 | Trima_12069 | 379 | 7.22e-31 | 75.6 |
