## Supplementary material for "Unusual genome expansion and transcription suppression in ectomycorrhizal *Tricholoma matsutake* by repetitive insertions of transposable elements": S2 Table

| Platform | Insert size (bp) | Number of reads | Average read length (bp) | Total bases (bp) |
| --- | --- | --- | --- | --- |
| HiSeq paired-end | 500 | 99,204,510 | 100 | 10,019,655,510 |
| HiSeq mate pair | 5,000 | 310,061,732 | 100 | 31,316,234,932 |
| MiSeq paired-end | 250 | 24,712,074 | 250 | 6,165,993,276 |

**S2 Table. The summary of genome sequencing**
