## Supplementary material for "Unusual genome expansion and transcription suppression in ectomycorrhizal *Tricholoma matsutake* by repetitive insertions of transposable elements": S3 Table

**S3 Table. Transposable element Pfam domains**

| Pfam ID | Pfam description |
| --- | --- |
| PF00075 | RNase H |
| PF00078 | Reverse transcriptase (RNA-dependent DNA polymerase) |
| PF00665 | Integrase core domain |
| PF02925 | Bacteriophage scaffolding protein D |
| PF02992 | Transposase family tnp2 |
| PF03184 | DDE superfamily endonuclease |
| PF03221 | Tc5 transposase DNA-binding domain |
| PF03732 | Retrotransposon gag protein |
| PF04687 | Microvirus H protein (pilot protein) |
| PF05699 | hAT family C-terminal dimerisation region |
| PF05840 | Bacteriophage replication gene A protein (GPA) |
| PF05970 | PIF1-like helicase |
| PF07727 | Reverse transcriptase (RNA-dependent DNA polymerase) |
| PF08283 | Geminivirus rep protein central domain |
| PF08284 | Retroviral aspartyl protease |
| PF10551 | MULE transposase domain |
| PF13358 | DDE superfamily endonuclease |
| PF13359 | DDE superfamily endonuclease |
| PF13456 | Reverse transcriptase-like |
| PF13837 | Myb/SANT-like DNA-binding domain |
| PF13976 | GAG-pre-integrase domain |
| PF14214 | Helitron helicase-like domain at N-terminus |
| PF14223 | gag-polypeptide of LTR copia-type |
| PF14529 | Endonuclease-reverse transcriptase |
