## Supplementary material for "Unusual genome expansion and transcription suppression in ectomycorrhizal *Tricholoma matsutake* by repetitive insertions of transposable elements": S4 Table

**S4 Table. Genomes used for comparative analysis**

| Species | Order | GenBank ID | Ecotype | Number of genes | Genome Size (Mbp) |
| --- | --- | --- | --- | --- | --- |
| *Agaricus bisporus* var. *bisporus* H97 | Agaricales | GCF_000300575.1 | Saprotroph | 10,448 | 30.2 |
| *Agaricus bisporus* var. *burnettii* JB137-S8 | Agaricales | GCF_000300555.1 | Saprotroph | 11,278 | 32.6 |
| *Amanita muscaria* Koide BX008 | Agaricales | GCA_000827485.1 | Ectomycorrhizal | 18,093 | 40.7 |
| *Auricularia subglabra* TFB-10046 SS5 | Auriculariales | GCF_000265015.1 | White-rot | 23,555 | 74.9 |
| *Botryobasidium botryosum* FD-172 SS1 | Cantharellales | GCA_000697705.1 | White-rot | 16,502 | 46.7 |
| *Coprinopsis cinerea* okayama7#130 | Agaricales | GCF_000182895.1 | Saprotroph | 13,356 | 36.2 |
| *Cylindrobasidium torrendii* FP15055 ss-10 | Agaricales | GCA_000934385.1 | White-rot | 13,936 | 31.6 |
| *Exidia glandulosa* HHB12029 | Auriculariales | GCA_001632375.1 | Saprotroph | 26,690 | 78.2 |
| *Fibularhizoctonia* sp. CBS 109695 | Atheliales | GCA_001630335.1 | Saprotroph | 32,854 | 95.1 |
| *Fistulina hepatica* ATCC 64428 | Agaricales | GCA_000934395.1 | Brown-rot | 11,244 | 33.8 |
| *Fomitiporia mediterranea* MF3/22 | Hymenochaetales | GCF_000271605.1 | White-rot | 11,338 | 63.4 |
| *Fomitopsis pinicola* FP-58527 SS1 | Polyporales | GCA_000344655.2 | Brown-rot | 13,852 | 41.6 |
| *Galerina marginata* CBS 339.88 | Agaricales | GCA_000697645.1 | White-rot | 21,391 | 59.4 |
| *Gloeophyllum trabeum* ATCC 11539 | Gloeophyllales | GCF_000344685.1 | Brown-rot | 11,755 | 37.2 |
| *Gymnopus luxurians* FD-317 M1 | Agaricales | GCA_000827265.1 | Saprotroph | 22,046 | 66.3 |
| *Hebeloma cylindrosporum* h7 | Agaricales | GCA_000827355.1 | Ectomycorrhizal | 15,376 | 38.2 |
| *Heterobasidion irregulare* TC 32-1 | Russulales | GCF_000320585.1 | Plant pathogen | 13,275 | 33.6 |
| *Hypholoma sublateritium* FD-334 SS-4 | Agaricales | GCA_000827495.1 | White-rot | 17,771 | 48 |
| *Jaapia argillacea* MUCL 33604 | Jaapiales | GCA_000697665.1 | White-rot | 16,375 | 45 |
| *Laccaria amethystina* LaAM-08-1 | Agaricales | GCA_000827195.1 | Ectomycorrhizal | 21,033 | 52.2 |
| *Laccaria bicolor* S238N-H82 | Agaricales | GCF_000143565.1 | Ectomycorrhizal | 18,215 | 64.9 |
| *Moniliophthora roreri* MCA 2997 | Agaricales | GCF_000488995.1 | Plant pathogen | 17,910 | 52.2 |
| *Neolentinus lepideus* HHB14362 ss-1 | Gloeophyllales | GCA_001632425.1 | Saprotroph | 13,157 | 35.6 |
| *Peniophora* sp. CONT | Russulales | GCA_001632445.1 | White-rot | 18,945 | 46 |
| *Piloderma croceum* F 1598 | Atheliales | GCA_000827315.1 | Ectomycorrhizal | 21,524 | 59.3 |
| *Pleurotus ostreatus* PC15 | Agaricales | GCA_000697685.1 | White-rot | 12,296 | 34.3 |
| *Plicaturopsis crispa* FD-325 SS-3 | Amylocorticiales | GCA_000827205.1 | White-rot | 13,617 | 34.5 |
| *Punctularia strigosozonata* HHB-11173 SS5 | Corticiales | GCF_000264995.1 | White-rot | 11,540 | 34.2 |
| *Pycnoporus coccineus* BRFM310 | Polyporales | GCA_002092935.1 | White-rot | 12,693 | 32.8 |
| *Rhizoctonia solani* AG-1 IA | Cantharellales | GCA_000334115.1 | Plant pathogen | 10,489 | 36.9 |
| *Schizophyllum commune* H4-8 | Agaricales | GCF_000143185.1 | White-rot | 13,194 | 38.5 |
| *Scleroderma citrinum* Foug A | Boletales | GCA_000827425.1 | Ectomycorrhizal | 20,995 | 56.1 |
| *Serendipita indica* DSM 11827 | Sebacinales | GCA_000313545.1 | Endophyte | 11,791 | 25 |
| *Serendipita vermifera* MAFF 305830 | Sebacinales | GCA_000827415.1 | Orchid mycorrhizal | 15,245 | 38.1 |
| *Sistotremastrum niveocremeum* HHB9708 | Trechisporales | GCA_001630475.1 | Saprotroph | 13,076 | 35.4 |
| *Sistotremastrum suecicum* HHB10207 ss-3 | Trechisporales | GCA_001632355.1 | Saprotroph | 13,654 | 33.9 |
| *Suillus luteus* UH-Slu-Lm8-n1 | Boletales | GCA_000827255.1 | Ectomycorrhizal | 18,303 | 37 |
