## Supplementary material for "Unusual genome expansion and transcription suppression in ectomycorrhizal *Tricholoma matsutake* by repetitive insertions of transposable elements": S5 Table

**S5 Table. Total bases and read counts for the three developmental stages of *Tricholoma matsutake***

| Development stage | Total bases | Read count | N (%) | GC (%) | Q20 (%) | Q30 (%) |
| --- | --- | --- | --- | --- | --- | --- |
| Hyphae | 10,954,625,640 | 108,461,640 | 0.004 | 49.58 | 94.12 | 86.92 |
| Primordia | 13,424,362,278 | 132,914,478 | 0.004 | 49.85 | 93.95 | 86.82 |
| Fruiting Body | 12,872,445,152 | 127,449,952 | 0.004 | 49.79 | 94.34 | 87.42 |
