## Supplementary material for "Unusual genome expansion and transcription suppression in ectomycorrhizal *Tricholoma matsutake* by repetitive insertions of transposable elements": S6 Table

**S6 Table. Target proteins responsible for genome defense by fungi**

| **Mechanism** | **Name** | **Description** | **Source organism** | **Length (aa)** | **GenBank accession** |
| --- | --- | --- | --- | --- | --- |
| Repeat-induced point mutation | Rid/Dim2^1^ | DNA methyltransferase | *Neurospora crassa* | 1454 | AAK49954.1 |
|  | Masc1^2^ | C5-DNA methyltransferase | *Ascobolus immersus* | 537 | AAC49849.1 |
|  | Masc2^3^ | C5-DNA methyltransferase | *Ascobolus immersus* | 1356 | AAC03766.1 |
| Meiotic silencing by unpaired DNA | Sad-1^4^ | RNA dependent RNA polymerase | *Neurospora crassa* | 1638 | EAA35012.3 |
|  | Sms-3^4^ | dicer-like protein | *Neurospora crassa* | 1584 | EAA32662.1 |
|  | Sms-2^4^ | meiotic silencing suppressor | *Neurospora crassa* | 989 | EAA29350.1 |
| Quelling | Qde-1^4^ | RNA-dependent RNA polymerase | *Neurospora crassa* | 1402 | EAA29811.1 |
|  | Dcl2^4^ | dicer-like protein | *Neurospora crassa* | 1396 | XP_963538.3 |
|  | Qde2^5^ | post-transcriptional gene silencing | *Neurospora crassa* | 938 | AAF43641.1 |

^1^Kouzminova, E. & Selker, E. U. dim-2 encodes a DNA methyltransferase responsible for all known cytosine methylation in *Neurospora*. The EMBO journal 20, 4309-4323, (2001).

^2^Malagnac, F. et al. A gene essential for de novo methylation and development in *Ascobolus* reveals a novel type of eukaryotic DNA methyltransferase structure. Cell 91, 281-290 (1997).

^3^Chernov, A. V., Vollmayr, P., Walter, J. & Trautner, T. A. Masc2, a C5-DNA-methyltransferase from *Ascobolus immersus* with similarity to methyltransferases of higher organisms. Biological chemistry 378, 1467-1473 (1997).

^4^Galagan, J. E. et al. The genome sequence of the filamentous fungus *Neurospora crassa*. Nature 422, 859-868, (2003).

^5^Catalanotto, C., Azzalin, G., Macino, G. & Cogoni, C. Gene silencing in worms and fungi. Nature 404, 245, (2000).
